# Reprogramming human B cells with custom heavy chain antibodies

**DOI:** 10.1101/2023.06.28.546944

**Authors:** Geoffrey L. Rogers, Chun Huang, Atishay Mathur, Xiaoli Huang, Hsu-Yu Chen, Kalya Stanten, Heidy Morales, Chan-Hua Chang, Eric J. Kezirian, Paula M. Cannon

## Abstract

We describe a genome editing strategy to reprogram the immunoglobulin heavy chain (IgH) locus of human B cells to express custom molecules that respond to immunization. These heavy chain antibodies (HCAbs) comprise a custom antigen-recognition domain linked to an Fc domain derived from the IgH locus and can be differentially spliced to express either B cell receptor (BCR) or secreted antibody isoforms. The HCAb editing platform is highly flexible, supporting antigen-binding domains based on both antibody and non-antibody components, and also allowing alterations in the Fc domain. Using HIV Env protein as a model antigen, we show that B cells edited to express anti-Env HCAbs support the regulated expression of both BCRs and antibodies, and respond to Env antigen in a tonsil organoid model of immunization. In this way, human B cells can be reprogrammed to produce customized therapeutic molecules with the potential for *in vivo* amplification.

## Introduction

Monoclonal antibodies are important therapeutics that allow specialized antibody designs not achieved by immunization, such as targeting self-antigens or highly variable pathogens.^1,2^ However, since their administration for chronic conditions can be burdensome and expensive, gene and cell therapies are also being considered for delivery.^3,4^ AAV vectors have been shown to support expression of custom antibodies from skeletal muscle,^4,5^ although such approaches have been hindered by low expression levels or immunogenicity in large animal^6,7^ and human studies.^8,9^ An alternative approach is to use genome editing to express custom antibodies from the natural immunoglobulin (Ig) locus in B cells.^10^ It is hypothesized that this would have the advantage of maintaining natural aspects of antibody production in the engineered cells, in particular the response to the matched antigen.

Several groups have now described such reprograming of the heavy chain locus (IgH) of mouse or human B cells.^11–18^ The complexity of IgH limits the sequences available for genome editing, but a common approach has been to target conserved sequences in the Eμ intron for insertion of a multicistronic cassette comprising both a custom V_H_ domain and a complete light (L) chain partner.^11–15^ Insertion at this site bypasses the endogenous V_H_ sequences and allows instead expression of an H chain comprising the custom V_H_ domain spliced to downstream endogenous constant regions, in addition to the L chain partner. B cells edited in this way appear to support the full range of B cell functions, including expression of membrane-bound B cell receptor (BCR) and secreted antibody isoforms, clonal expansion, anamnestic responsiveness, class switch recombination, and somatic hypermutation.^11,12^

The V_H_ plus L chain insertion approach presents certain challenges. It requires expression of both H and L chain sequences from the same cassette, which can be achieved by including a self-cleaving peptide^11,12,14^ or a long flexible linker.^11,13^ More significantly, edited cells could still express endogenous H and L chains from unedited loci that could mispair with the engineered antibody chains and create unwanted antigen specificities, although a separate editing step to disrupt at least the endogenous Igκ L chain can also be included.^11,14^ Finally, since the Fc component of the engineered antibody is determined by the endogenous IgH sequences, it is not possible to customize this component to include beneficial modifications.^19,20^

We report here an alternate and simplified approach to engineering human B cells. It is based on the insertion of a monocistronic cassette, comprising a B cell specific promoter and an antigen recognition domain, downstream of the CH1 exon in IgH constant regions. By excluding CH1, we mimicked the design of H chain antibodies (HCAbs) found in camelids, which do not have L chain partners and derive their antigen specificity from a single V_H_H domain.^21,22^ The HCAb editing approach supports the use of a variety of different molecules to serve as antigen recognition domains, including both antibody (V_H_H, scFvs) and non-antibody components. By inserting anti-HIV Env recognition domains in IgG1, we confirmed that HCAb-engineered B cells exhibit physiologically regulated expression of both BCR and secreted HCAb isoforms and respond to immunization with the specific antigen in a tonsil organoid model. Finally, we demonstrated the additional flexibility of this approach by targeting an alternate site within the constant region, which will also allow customization of Fc sequences.

## Results

### Genome editing strategy to produce human H chain only antibodies

Reprogramming B cells to express custom antibodies is challenging due to the complex design of the Ig locus. Human antibodies are encoded across three loci (IgH for H chains and Igκ or Igλ for L chains) and undergo sequence rearrangements to produce the V_H_ and V_L_ variable domains that determine the antigen specificity of each B cell clone. In addition, class switch recombination events in IgH can pair V_H_ sequences with alternate constant domains that provide the Fc sequences of the antibody. The resulting heterogeneity of genomic sequences in individual B cell clones therefore limits the number of conserved sites that could be targeted for genome engineering.

We addressed this problem by mimicking the design of camelid H chain antibodies (HCAbs). These antibodies do not pair with L chains since they lack the CH1 domain that is the major determinant of H-L interactions. We achieved a similar design by insertion of single polypeptide antigen recognition domains (ARDs) into the intron downstream of CH1 (**Fig. 1a**). The inserted cassettes comprised a B cell specific promoter driving expression of the ARD, and a splice donor to create chimeric transcripts that link the ARD to the rest of the endogenous constant exons. We hypothesized that engineered HCAbs would retain the ability to express both BCR and secreted antibody isoforms, and that this would be controlled by the elements of the Ig locus that regulate this alternative splicing.^23^ In this way, the edited B cells should secrete the custom HCAb in response to antigen, while avoiding the possibility that the engineered H chain would mispair with endogenous L chains to create unwanted antigen specificities.

**Figure 1.**
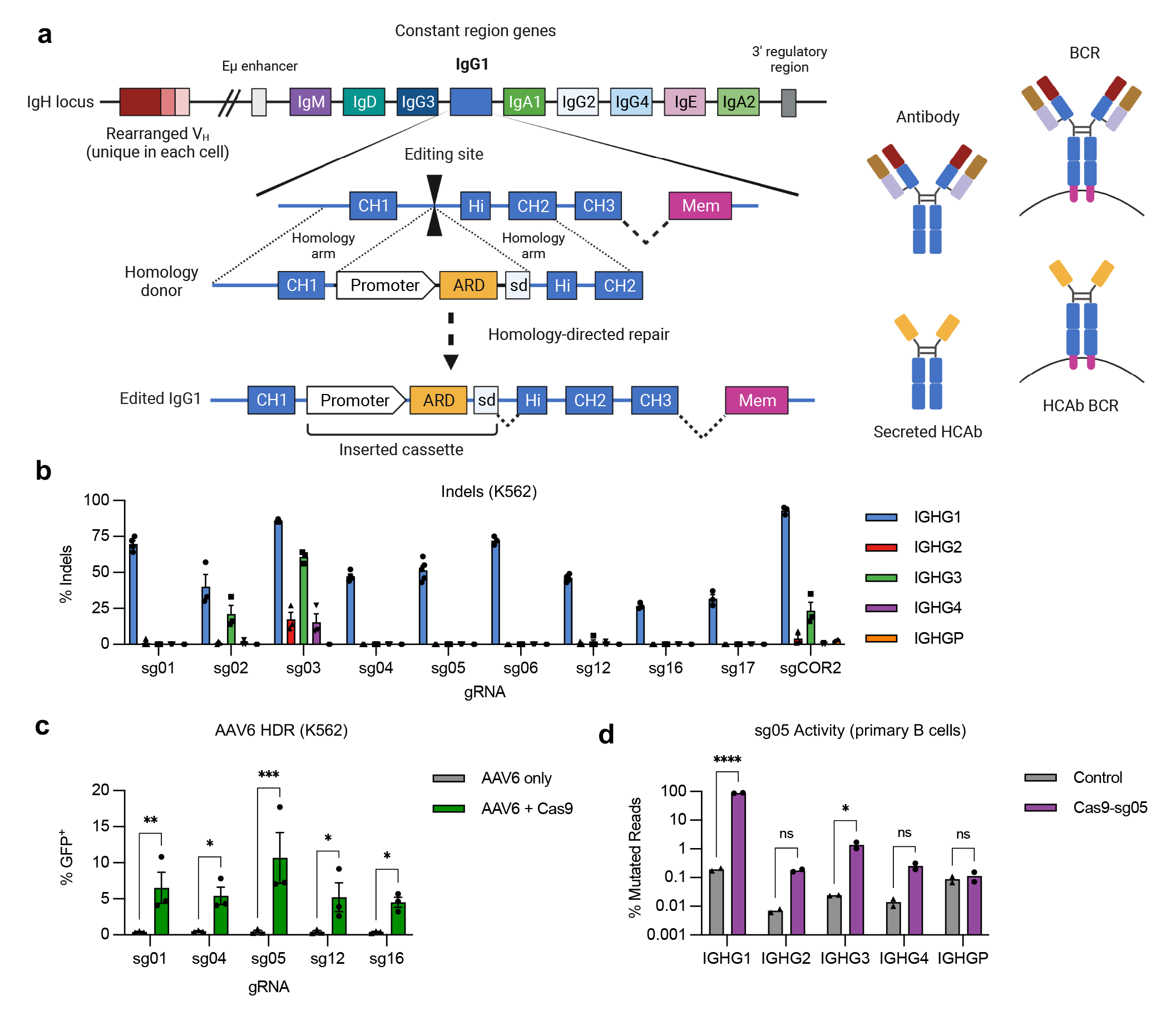
Genome editing at the constant region of the IgH locus. (**a**) Antibody H chains are encoded by a rearranged variable domain (V_H_), spliced to a constant region that can be altered by class switch recombination. Alternate splicing within the constant region generates secreted antibody or membrane anchored (BCR) isoforms. Genome editing to insert custom antigen recognition domains (ARD) downstream of CH1 exons in the IgH constant regions can create Heavy chain only antibodies (HCAbs). The example shown is targeting human IgG1, with the expanded view showing the constant region exons. Genome editing is directed by homology donors containing 500-750 bp homology arms, flanking CRISPR/Cas9 target sites in the intron downstream of CH1. Homology directed repair of the targeted DNA break inserts a cassette comprising a B cell-specific promoter, a custom ARD, and a splice donor to direct splicing to downstream endogenous exons. (**b**) K562 cells were edited with Cas9 RNPs programmed by gRNAs targeting the IgG1 CH1 intron and indels measured at on-target (IGHG1) and off target (IGHG2-P) genes by Sanger sequencing and ICE (*n* = 3). (**c**) K562 cells were edited by Cas9 RNPs plus AAV6 homology donors containing a GFP expression cassette, matched for each gRNA. GFP expression was measured after 3 weeks, to dilute out episomal AAV genomes (*n* = 3). (**d**) On-and off-target activity of sg05 was measured at indicated IGHG genes in primary human B cells (*n* = 2), 5 days after editing with sg05 RNPs, by targeted amplicon deep sequencing, with percentage mutated reads calculated as insertions, deletions, ≥ 2 bp changed. See also Extended Data Fig. 1d,e. Error bars show mean ± SEM. Statistical comparisons (c-d) were performed by 2-way ANOVA. * *p* < 0.05, ** *p* < 0.01, *** *p* < 0.001, **** *p* < 0.0001.

We used CRISPR/Cas9 gene editing to target the 391bp intron between the CH1 and hinge exons of IgG1. The high degree of homology (92-96%) of this intron with sequences in IgG2-4, and the pseudogene IgGP, limited the potential target sequences (**Fig. S1**). Using a panel of 10 *S. pyogenes* Cas9 guide RNAs (**Supplementary Table 1**), we first evaluated on-and off-target cutting activity in a cell line screen with limited sensitivity (**Fig. 1b**), which eliminated 3 candidates. We then evaluated the remaining gRNAs for the ability to support site-specific insertion at the targeted IgG1 sites by homology-directed repair (HDR), using single-stranded oligonucleotide (ssODN), plasmid, or AAV6 vectors as homology donors (**Fig. 1c, Extended Data Fig. 1a,b**). These analyses identified gRNA sg05 as supporting the highest HDR editing rates, with precise insertion confirmed by in-out PCR (**Extended Data Fig. 1c**) and Sanger sequencing (**Supplementary Fig. 2**). A more in-depth analysis of sg05 activity in primary B cells showed some off-target activity at the IGHG3 gene (1-1.7%) and low amounts at IGHG2 and IGHG4 (0.16- 0.32%), whereas activity at the on-target IGHG1 site exceeded 86% (**Fig. 1d, Extended Data Fig. 1d,e**). These data therefore supported the selection of sg05 for studies of HCAb editing in human B cells.

### HCAb expression and activity in edited B cell lines

We evaluated the HCAb engineering strategy in human B cell lines using ARDs based on the J3^24^ and A6^25^ camelid V_H_H domains, both of which recognize the HIV-1 gp120 Env protein. Plasmid homology donors, compatible with the sg05 target site, were co-electroporated into Raji B cells with sg05 Cas9 RNPs. Specific HCAb-BCR expression was evaluated by flow cytometry for surface IgG, which is not normally expressed by Raji cells,^26^ and by binding to recombinant gp120. Cells receiving both RNPs and homology donors had higher levels of surface staining than control cells or cells receiving the homology donors alone, and the correlation between IgG and gp120 staining suggested both labels were binding the same BCR molecule (**Fig. 2a**). The relatively low editing rates we obtained reflect the inefficiency of plasmid homology donors in the Raji cell line, and similar editing rates were achieved with donors containing a control GFP expression cassette, or in the alternate Ramos B cell line (**Extended Data Fig. 2a-b**).

**Figure 2.**
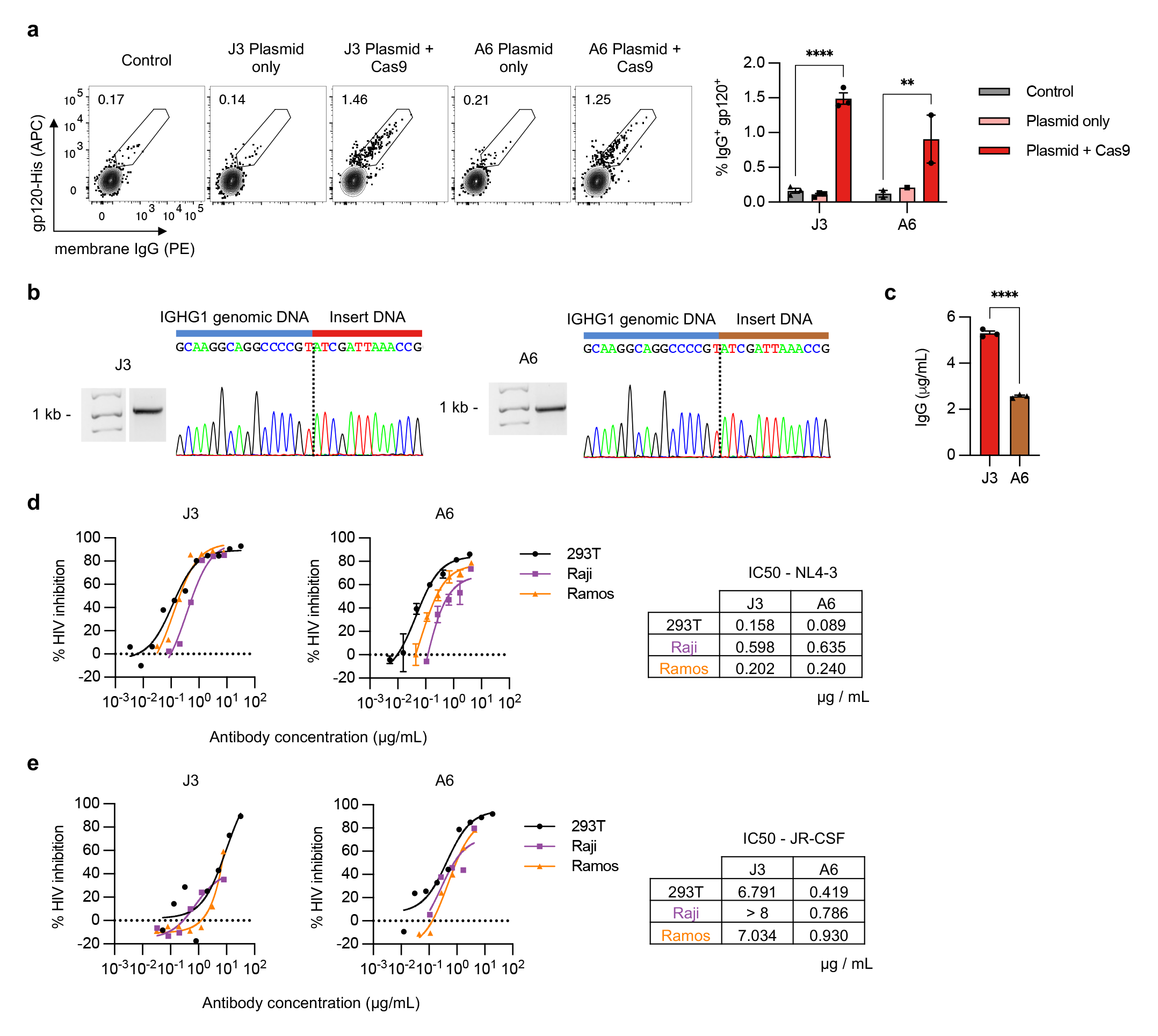
Engineering B cell lines to express anti-HIV HCAbs. (**a**) Raji B cells were edited with sg05 Cas9 RNPs plus plasmid homology donors encoding J3 (*n* = 3) or A6 (*n* = 1-2) V_H_H cassettes. Surface expression of resulting HCAb BCRs was measured 1 week later by flow cytometry, staining for surface IgG expression and gp120 binding. Representative plots are shown, together with summary data. (**b**) J3 or A6 edited Raji cells were FACS-sorted based on surface IgG (see Extended Data Fig. 2c), and the enriched population was subjected to in-out PCR and Sanger sequencing of PCR bands to confirm precise insertions. The dotted line indicates the predicted sg05 cut site. Uncropped gel is available in Supplementary Fig. 3b. (**c**) Sorted J3- or A6-edited Raji cells were seeded at 10^6^ cells/mL and IgG secretion measured 2 days later by ELISA (*n* = 3). (**d**) J3 or A6 HCAbs (*n* = 3 technical replicates) from supernatants of sorted populations of edited Raji and Ramos cells were analyzed for anti-HIV neutralization activity against X4-tropic HIV strain NL4-3 using the TZM-bl assay. Control recombinant HCAb supernatants were obtained from transfected 293T cells. Neutralization curves are shown for serial dilutions of each HCAb and IC50s were calculated from the curves. (**e**) Anti-HIV neutralization activity determined as in (d), against R5-tropic HIV strain JR-CSF. Error bars show mean ± SEM. Statistics in panel (a) were performed by 2-way ANOVA. ** *p* < 0.01, **** *p* < 0.0001.

To further evaluate HCAb editing, edited Raji and Ramos cells were sorted based on expression of GFP or surface IgG, as appropriate (**Extended Data Fig. 2c-d**). Specific in-out PCR and Sanger sequencing confirmed site-specific insertion for all 3 inserts in Raji cells **(Fig. 2b, Extended Data Fig. 2e**). Since these cell lines do not secrete human IgG,^26,27^ HCAb secretion could be measured using an anti-IgG ELISA. Secreted IgG was detected in the supernatants of J3 or A6 edited cells, but not GFP-edited cells (**Fig. 2c, Extended Data Fig. 2f**). Together these data support that the chimeric HCAb transcripts arising from the inserted cassettes could be alternatively spliced to produce both membrane-bound and secreted isoforms.

Finally, the functionality of the secreted HCAbs was evaluated. Raji cells seeded in equal numbers secreted higher levels of J3 than A6 HCAbs (**Fig. 2c**). When normalized, we observed similar HIV neutralization activities for HCAbs originating from either edited Raji or Ramos cells, or the matched recombinant HCAbs produced from transfected 293T cells (**Fig 2d,e**). Moving forward, we chose to focus on J3 due to its higher secretion levels and superior anti-HIV potency across broad panels of HIV strains.^25,28^

We next tested the hypothesis that the HCAb design would bypass the concern of H and L chain mispairing. We co-expressed the J3 HCAb in 293T cells with both the H and L chains of a conventional monoclonal human antibody, or its L chain only, and evaluated interactions using ELISAs and antibodies specific for V_H_H domains or human L chains (**Extended Data Fig. 3**). We did not observe any such interactions for the HCAb/L chain combinations. In contrast, we did detect the expected H chain heterodimers when the HCAb was co-expressed with the H+L complete antibody combination. However, such pairings are expected to create bispecific antibodies that keep intact each separate antigen recognition domain, limiting the possibilities for novel specificities or self-reactivity. Together, these data support the idea that HCAb expression in a B cell that also expresses endogenous H and L chains will provide a better safety profile than alternate B cell editing approaches that result in co-expression of both engineered and endogenous H and L chains.

**Figure 3.**
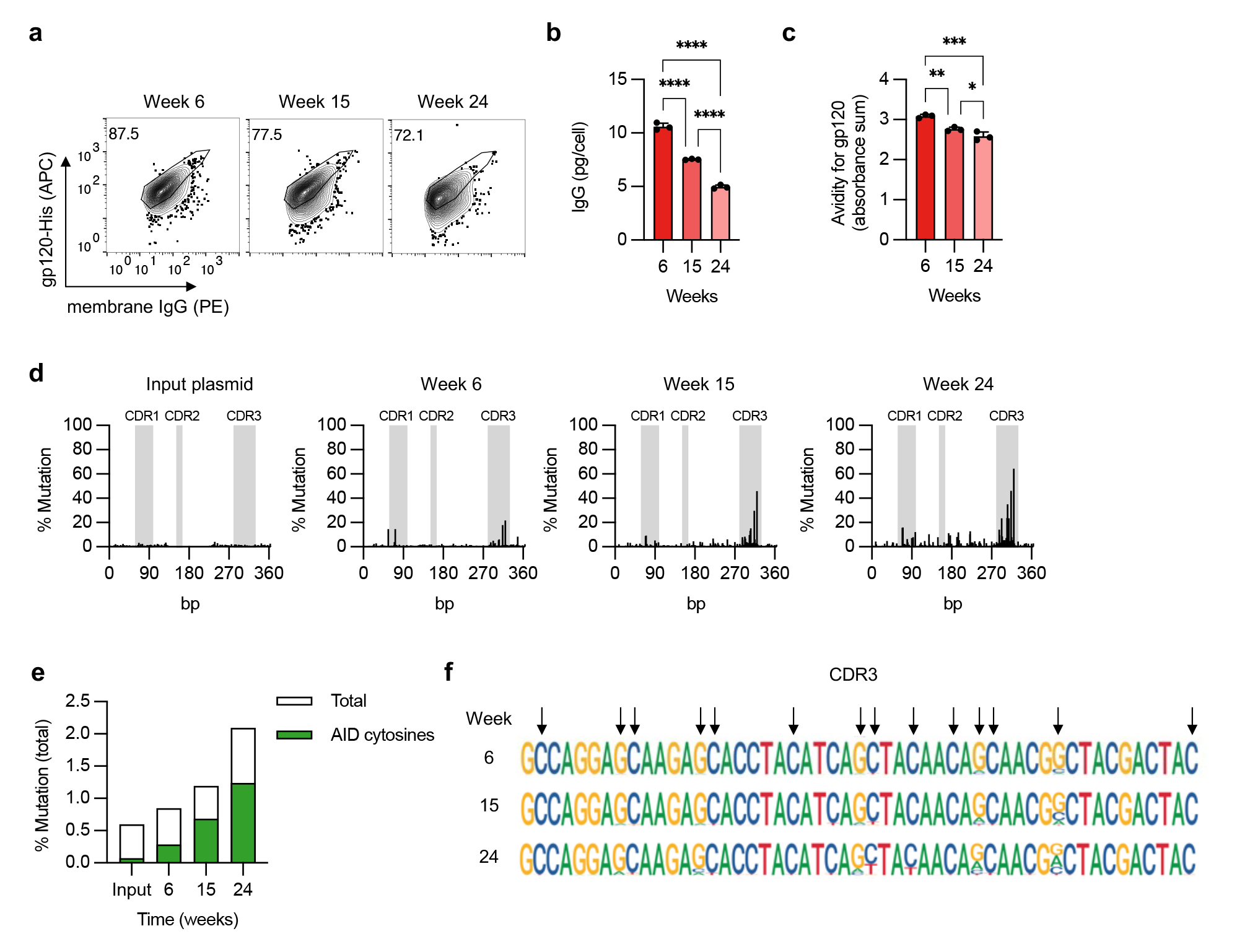
Evidence for somatic hypermutation of inserted J3 V_H_H sequences. (**a-d**) FACS-sorted J3-edited Raji cells were cultured for 6 months without further selection. (**a**) J3-BCR expression by flow cytometry at indicated times, gated on IgG^+^ cells. (**b**) Cells from each time point were seeded at 10^6^ cells/mL for 2 days and IgG secretion measured by ELISA, normalized for number of cells seeded. (**c**) Supernatants were assessed for gp120 binding by ELISA, across a range of normalized IgG concentrations. Absorbances across the curve were summed as a surrogate for area under the curve. See also Extended Data Fig. 4a. (**d**) Changes at J3 sequence in edited cells over time, measured by deep sequencing and compared to input homology donor plasmid. Percentage mutation at each position was calculated as the frequency of reads that did not match the wild-type sequence. CDR regions are indicated in gray. (**e**) Mutations identified by deep sequencing of J3-edited cells at each timepoint were summed and divided by the total sequence length to determine total % mutations. Shown in green are mutations associated with AID hotspot motif cytosines (WR<u>C</u>H). (**f**) Sequence logo plot of the CDR3 region of J3 at each time point. Arrows indicate AID hotspot cytosines, on either strand. Error bars show mean ± SEM. Statistics in panel (b-c) were performed by 1-way ANOVA.

### Evidence for somatic hypermutation in HCAb-edited B cell lines

Somatic hypermutation (SHM) is a natural evolutionary process in B cells driven by the enzyme activation-induced cytosine deaminase (AID) that alters variable domain sequences in both H and L chains and allows antibody affinity maturation.^29^ Although the mechanism whereby AID targets antibody variable domains is complex and not well-understood, the link to transcription start sites in the Ig locus^30^ suggested that promoter-driven expression of inserted ARDs in HCAb-edited B cells could still support SHM, despite being inserted in a constant region (**Fig 1a**).

To evaluate this possibility, we cultured J3-edited Raji cells that constitutively express AID^31^ for 6 months without selection. We looked for evidence of SHM by deep sequencing at different timepoints, and for changes in gp120 binding that could reflect the acquisition of mutations. Over time, there was a progressive loss of gp120 binding for both the BCR and secreted J3 isoforms, along with a reduction in J3 HCAb secretion, suggesting mutations in the J3 sequence that altered functionality and expression (**Fig. 3a-c, Extended Data Fig. 4a**). Deep sequencing confirmed that the J3 sequence in Raji cells accumulated mutations over time, and in particular within the CDR3 sequence that is generally considered to be the most important region for antigen binding (**Fig. 3d**). The observed mutations were predominantly at cytosine residues within predicted AID hotspots (**Fig. 3e,f**), consistent with AID-mediated mutagenesis.

**Figure 4.**
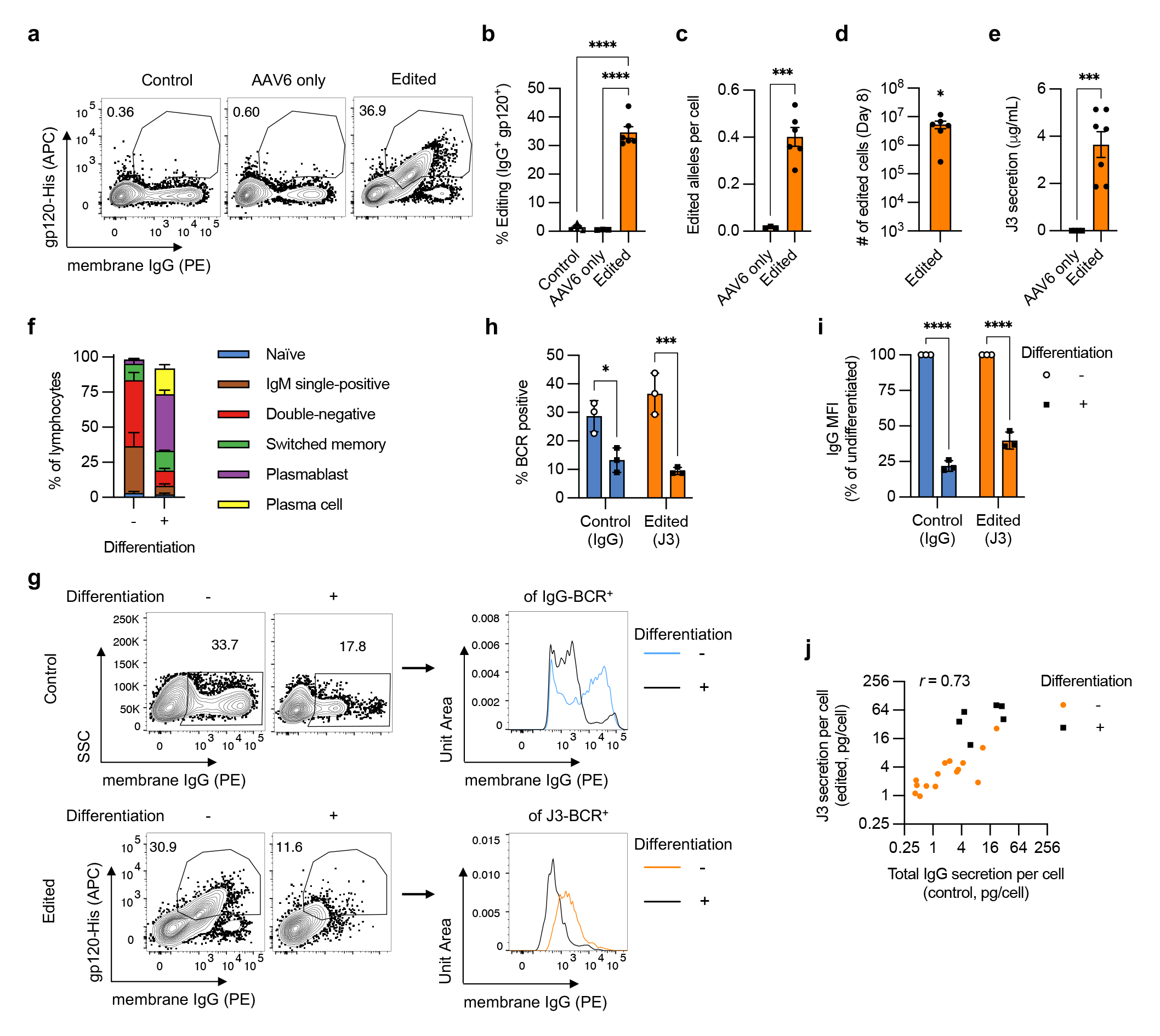
Engineering primary human B cells. Primary human B cells from *n* = 3-6 independent experiments were activated with BAC in XF media, starting at day -3, and edited at day 0 with sg05 Cas9 RNPs and an AAV6- J3 homology donor (MOI = 5 x 10^5^ vg/cell). (**a-b**) Editing rates were measured at day 8 by flow cytometry for surface J3-BCR. (**c**) Editing was quantified by in-out ddPCR at day 8 and normalized per cell against a control reaction. (**d**) The yield of edited cells at day 8 was calculated from 5 x 10^5^ starting B cells. (**e**) J3 HCAb secretion was measured by gp120-IgG ELISA at day 8. (**f-h**) Edited B cells in BAC culture were differentiated by switching to the DP protocol on day 2 post-editing and cultured for a further 6 days. Undifferentiated populations were maintained in BAC until day 8. (**f**) B cell phenotypes, measured on day 8 by flow cytometry, as described in Supplementary Fig. 5. (**g-i**) Expression of total IgG-BCR (control, unedited samples) and J3-BCR (edited samples) was measured by flow cytometry at day 8 for cells without or with differentiation *(n* = 3). The intensity of membrane IgG expression was evaluated for IgG^+^ control B cells and J3^+^ edited B cells. Shown are representative plots (**g**) and summary graphs of IgG-or J3-BCR positivity (**h**) and expression levels (**i**) after differentiation, normalized to paired undifferentiated samples. Data for edited (J3) but undifferentiated cells in panel (h) is a subset of the data in panel (b). (**j**) Scatter plot comparing the rate of total IgG secretion from control unedited samples versus J3 HCAb secretion from edited samples. Each point represents measurements from paired samples from the same experiment. Samples were collected at several time points after editing, and differentiated samples are also identified. The Pearson correlation is indicated. Error bars show mean ± SEM. Statistics were calculated by 1-way ANOVA (b), 2-tailed t-test (c,e), 1- sample t-test (d), or 2-way ANOVA (h-i). * *p* < 0.05, *** *p* < 0.001, **** *p* < 0.0001.

Interestingly, our data suggests that AID activity was also influenced by the specific sequence of the inserted DNA cassette. Raji cells containing a control GFP cassette did not develop such a mutational signature, and neither did the A6 V_H_H domain (**Extended Data Fig. 4b,c**). This was not due to a lack of AID hotspot motifs, as these were present in all 3 sequences evaluated (**Extended Data Fig. 4d,e**), suggesting that additional features of the antibody sequence beyond AID hotspots are needed to license SHM activity.

### Activation and culture of primary human B cells

Optimizing cell culture conditions is critical for successful *ex vivo* cell engineering. For B cell genome editing, we focused on three major criteria. First, we wanted to promote cell growth and cell cycling, since this is necessary for HDR genome editing.^32^ Second, an ability to differentiate the edited cells towards antibody-secreting cells (ASCs), which would allow us to evaluate both HCAb-BCR expression in the initially edited cells and HCAb secretion from the resulting ASCs. Finally, we wanted to establish edited B cells with memory phenotypes. Such cells can respond rapidly to antigen, which is expected to drive expansion and HCAb secretion in a therapeutic *in vivo* application.

We evaluated 3 different B cell stimulation conditions: a 3-step differentiation protocol (DP) we and others have previously used for human B cell genome editing,^33,34^ an anti-RP105 antibody,^11,14^ and a commercial B cell activation cocktail (BAC) (**Supplementary Fig. 4**). Each protocol was assessed in 3 different basal media for cell expansion, viability, cell size and IgG secretion (**Extended Data Fig. 5a-e**). As expected, the DP treatment resulted in little expansion of cell numbers but robust ASC differentiation and IgG secretion. Unexpectedly, in our hands anti-RP105 did not support either B cell survival or expansion. In contrast, BAC stimulation in XF media drove robust >100-fold B cell expansion, with or without FBS.

**Figure 5.**
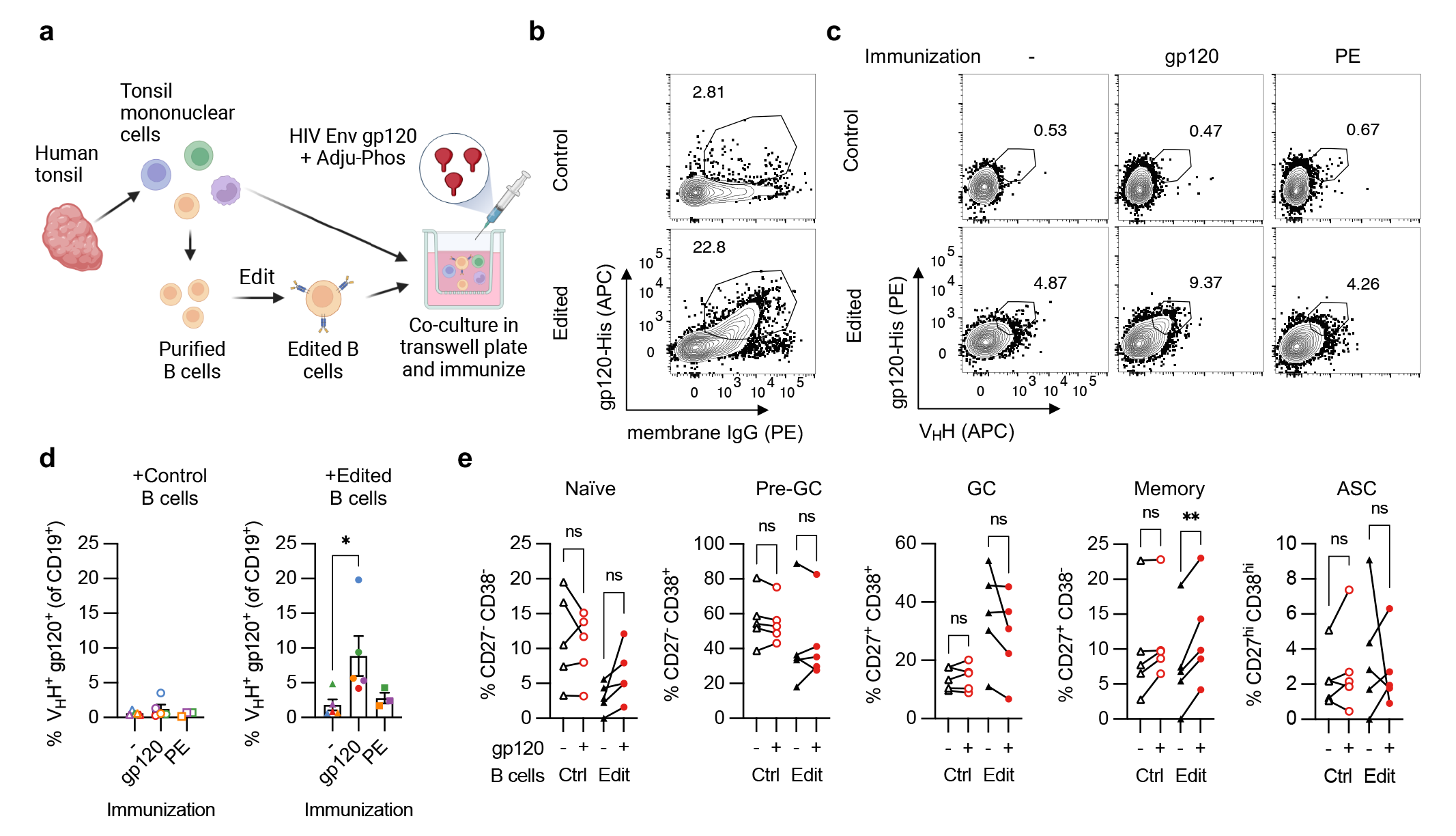
Antigen-specific expansion of HCAb-engineered B cells. (**a**) Diagram of tonsil organoid system. (**b**) Representative plots of control or J3-edited tonsil B cells, treated with RNPs plus AAV6-J3 at MOI = 5 x 10^5^ vg/cell. (**c-e**) Tonsil B cells from *n* = 3-5 donors were edited with RNPs plus AAV6-J3 at MOI = 1-5 x 10^5^ vg/cell, cultured for 2-4 days, and reconstituted with total autologous TMNCs. Cultures were immunized with HIV gp120 plus Adju-Phos or control PE protein plus Alhydrogel and cells were harvested 12 days later for analyses. (**c**) Representative panel showing increase in gp120^+^ V_H_H^+^ cells after immunization with gp120 but not PE, measured by flow cytometry, gated on CD19^+^ CD3^-^ B cells as described in Supplementary Fig. 8. (**d**) Response of tonsil organoid cultures containing control or edited B cells to gp120 or PE immunization. Matching colors indicate samples from the same tonsil donor. (**e**) Phenotypes of B cells in tonsil organoids containing control or J3-edited B cells were characterized at day 12 by flow cytometry, with or without gp120 immunization, using the gating strategy described in Supplementary Fig. 8. Error bars show mean ± SEM. Statistics in panel (d-e) were calculated by 1-way ANOVA. * *p* < 0.05, ** *p* < 0.01.

We further analyzed the phenotypes of the B cells in the different cultures. We found that BAC culture drove activation and class switching from naïve to IgM single-positive cells, double-negative cells (likely memory or ASC precursors),^35^ and memory B cells (**Extended Data Fig. 5f**). Further, cells initially activated with BAC could still be differentiated towards ASCs upon subsequent DP treatment (**Extended Data Fig. 5f**), with IgG secretion capacity comparable to cells undergoing only the DP protocol (**Extended Data Fig. 5g**). We therefore selected BAC activation in serum-free XF media as the protocol of choice, since B cells treated in this way both expanded in culture and avoided terminal ASC differentiation, while also retaining competence for further differentiation into ASCs under defined conditions.

### HCAb editing in primary human B cells

To edit primary B cells, we adapted our previously published protocol using AAV6 vectors to deliver homology donor DNA.^33^ AAV6-J3 donors were combined with sg05 RNPs and first evaluated on the Raji and Ramos B cell lines, where we observed up to 10-fold increased editing rates compared to those achieved with matched plasmid donors (**Extended Data Fig. 6a**). With primary B cells, AAV6-J3 donors supported editing rates of approximately 35%, measured by flow cytometry for surface J3-BCR expression, or ddPCR for edited IGHG1 alleles (**Fig. 4a-c**). Site-specific insertion of the J3 cassette was also confirmed by in-out PCR and Sanger sequencing (**Extended Data Fig. 6b**). The edited primary B cells retained the ability to undergo robust expansion after editing (**Fig. 4d**), and secreted J3 HCAbs (**Fig. 4e**). The engineered antibodies also had the expected ability to neutralize HIV-1 strain NL4-3 (**Extended Data Fig. 6c**)

**Figure 6.**
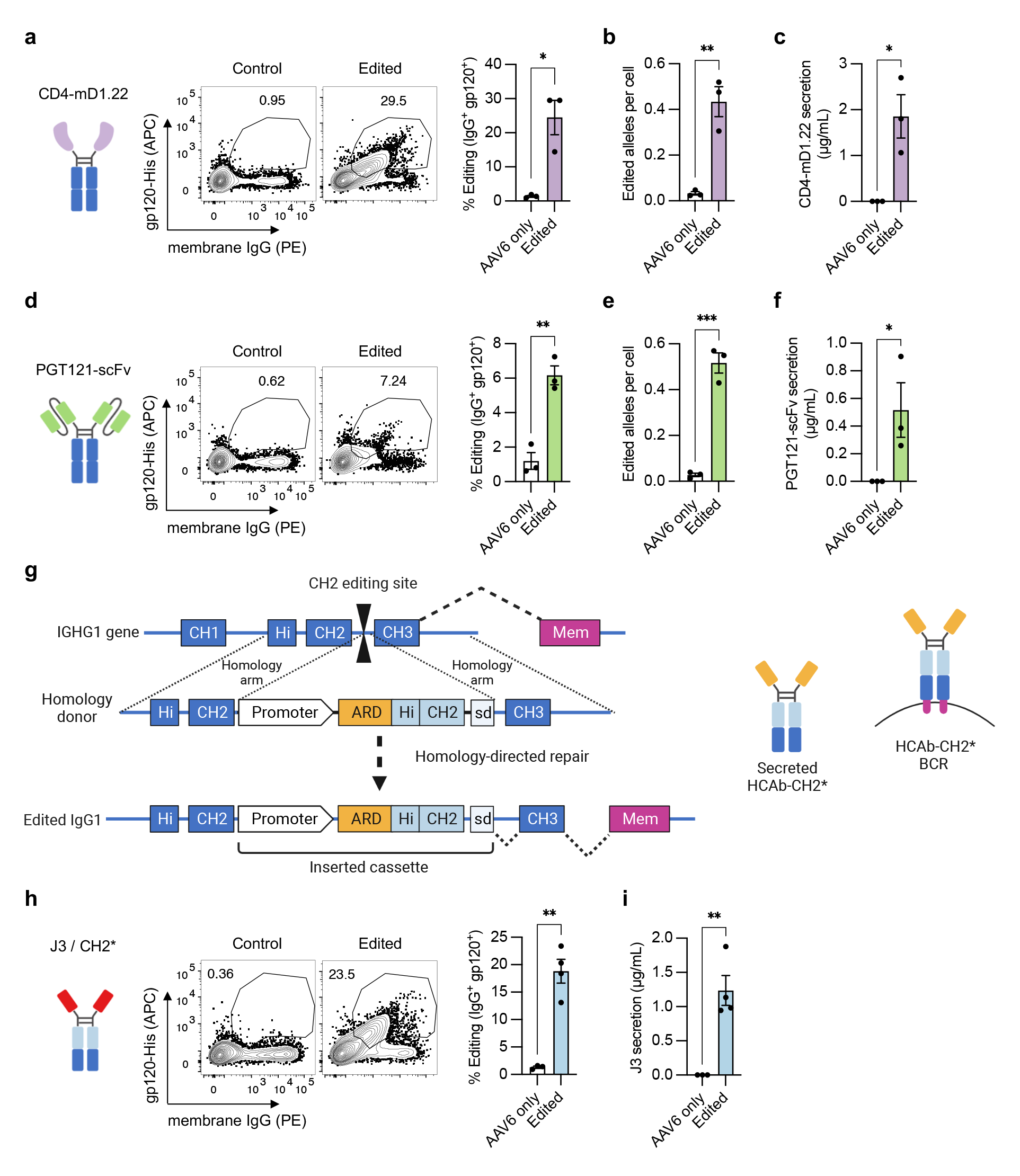
Editing with alternative HCAb structures and modified Fc domains. (**a-f**) Primary human B cells from *n* = 3 independent experiments were activated with BAC in XF media, starting at day -3, and edited at day 0 with sg05 Cas9 RNPs and AAV6-CD4 donors (a-c, MOI = 5 x 10^5^ vg/cell) or AAV6-PGT121-scFv donors (d-f, MOI = 5 x 10^5^ vg/cell). (**a,d**) Editing rates were measured at day 8 by flow cytometry for surface J3-BCR. (**b,e**) Editing was quantified by in-out ddPCR at day 8 and normalized per cell against a control reaction. (**c,f**) HCAb secretion was measured by gp120-IgG ELISA at day 8. (**g**) Design of CH2 editing approach. Cas9 gRNA CH2-g1 targets the intron downstream of CH2. Homology donor cassette contains B cell specific promoter, antigen recognition domain (ARD), codon-wobbled IgG1 Hinge (Hi) and CH2 exons, and splice donor to link to endogenous CH3 and membrane exons after insertion. (**h-i**) Primary human B cells from *n* = 3-4 independent experiments were activated with BAC in XF media, starting at day -3, and edited at day 0 with CH2-g1 Cas9 RNPs and AAV6-J3-CH2* donor (MOI = 10^4^ vg/cell). **(h)** Editing rates were measured at day 8 by flow cytometry for surface J3-BCR. (**i**) J3 HCAb secretion was measured by gp120-IgG ELISA at day 8. Error bars show mean ± SEM. Statistics were calculated by 2-tailed t-test. * *p* < 0.05, ** *p* < 0.01, *** *p* < 0.001.

By varying the AAV6-J3 MOIs, we observed that although lower MOIs reduced the initial editing rates achieved, they also had less of an impact on cell proliferation, resulting in similar total numbers of edited cells by day 8 (**Extended Data Fig 6d-h**). The inhibition of proliferation at higher doses could be due to the reported p53-mediated sensing of AAV ITRs.^36^ Editing was also possible with all of the BAC and DP activation protocols we evaluated, although none outperformed BAC in serum-free XF media (**Supplementary Fig. 6**).

We next investigated the differentiation capability of edited B cells under different culture conditions. The editing process did not significantly impact the phenotype of BAC cultured cells, which remained largely IgM single-positive, double-negative, and memory B cells (**Fig. 4f, Extended Data Fig. 6i**). After editing, the cells could also still be induced towards ASCs (plasmablasts and plasma cells), with some enhancement of plasmablast generation compared to unedited populations (**Fig. 4f, Extended Data Fig. 6i**).

When B cells differentiate towards ASCs, there is a switch from BCR to secreted antibody isoforms.^23^ After differentiation of unedited B cells we observed around a 50% reduction in the frequency of IgG-BCR^+^ cells and 75% reduction in the amount of IgG-BCR on the cell surface (**Fig. 4g-i**). Expression of J3-BCR on edited B cells paralleled these changes, with similar reductions in both the frequency and magnitude of J3-BCR expression after differentiation (**Fig. 4g-i**). The switch from J3-BCR to antibody isoforms was also confirmed at the mRNA level (**Extended Data Fig. 6j**). We also observed a similar concordance for the secretion of antibodies, since the rate of IgG secretion per cell in unedited samples correlated with the rate of J3 secretion per cell from edited samples, across different matched time points and differentiation conditions (**Fig. 4j**). As expected, both total IgG and J3 HCAb secretion rates per cell were generally highest in the differentiated samples.

In sum, these data show that primary human B cells activated with BAC can be efficiently edited at IGHG1. HCAb editing has minimal impact on B cell differentiation, with the cells adopting a memory or precursor phenotype. Finally, the edited cells can undergo differentiation, which enhances antibody secretion and reduces BCR expression. These changes were similar for endogenous IgG and J3 HCAb, implying proper regulation of this important process for the HCAb construct.

### HCAb-engineered B cells respond to immunization in tonsil organoids

We used a tonsil organoid system^37^ to examine the ability of HCAb-engineered human B cells to respond to stimulation by a matched antigen (**Fig. 5a**). We first confirmed the ability of non-engineered tonsil B cells to respond to immunization in this model by treating the organoid cultures with HIV gp120 or PE antigens, and further identified an effective adjuvant for gp120 (**Supplementary Fig. 7**). Next, B cells were purified from total tonsil mononuclear cells (TMNCs) and edited as before using sg05 RNPs and AAV6-J3 vectors (**Fig. 5b**). Four days later, the edited B cell population was reconstituted with total TMNCs (including an excess of unmanipulated B cells) and paired samples were cultured without immunization, or immunized with gp120 or control PE protein plus adjuvant. Immunization with gp120 resulted in a significant expansion of J3-edited B cells compared to the unimmunized cultures (**Fig. 5c,d**). No expansion was observed following immunization with PE, confirming that this was an antigen-specific response. Finally, phenotyping of B cells demonstrated that memory phenotypes developed specifically in the J3-edited cells after gp120 immunization (**Fig. 5e**).

### Flexibility of the HCAb editing platform

The simplicity of the HCAb design and editing strategy suggests that other antibody derivatives, and even non-antibody components, could be used to mimic antigen recognition and drive B cell responses. To evaluate this, we engineered peripheral blood B cells with two alternative ARDs that also recognize HIV gp120: a stabilized variant of domain 1 of CD4, the natural receptor for HIV (CD4-mD1.22),^38^ and an scFv derived from the anti-HIV broadly neutralizing antibody PGT121.^39^ Cell surface expression and antibody secretion were confirmed by flow cytometry and ELISA, respectively, and editing was confirmed by ddPCR (**Fig. 6a-f**). Despite comparable editing rates at the DNA level, we observed lower surface expression and secretion of the PGT121 scFv HCAb than for CD4-mD1.22, or our previous data with J3. This may indicate that this scFv requires further optimization for expression and functionality, as is common for such derivatives.^40^

We next hypothesized that using alternate editing sites within IgH constant region domains would allow us to also reprogram components of the HCAb beyond the ARD. This would be useful for applications requiring modification of an antibody’s effector functions or protein stability, which are largely determined by the Fc region.^19,20^ To test this possibility, we developed genome editing tools targeting the intron downstream of IGHG1 CH2 (**Supplementary Table 1**). The inserted cassette comprised the B cell specific promoter and J3 ARD as before, but additionally contained sequences encoding the IgG1 Hinge and CH2 exons. These would replace the endogenous sequences in the resulting HCAb, and a splice donor links the insert to the remaining CH3 and membrane exons of IGHG1 (**Fig. 6g**).

A modification to the designed homology donor was necessary to enable this alternative insertion site, since homology between the donor and endogenous hinge-CH2 sequences was expected to interfere with the desired homology-directed edit. This was achieved by codon wobbling the hinge-CH2 sequences in the insert to reduce homology with the endogenous sequences. Primary B cells edited by this approach showed the expected J3 HCAb surface expression and antibody secretion (**Fig. 6h-i**), validating the functionality of this approach. Together, these results demonstrate the flexibility of the HCAb editing platform, allowing customization of both antigen recognition and Fc domains.

## Discussion

The therapeutic success of reprogrammed CAR T cells has generated interest in the possibility of engineering other immune cells as cellular therapies.^10,41^ For B cells, an ability to customize the antibodies that are produced would allow expression of specific therapeutic antibodies, including those that cannot be generated naturally through vaccination. Other applications of engineered B cells could take advantage of the secretory potential of differentiated plasma cells, which produce up to 10,000 antibodies per cell per second^42^ and can persist for years in a terminally differentiated quiescent state.^43^ They therefore have the potential to act as *in vivo* factories, secreting both antibodies and non-antibody proteins.^34,44^ B cells also form memory cells, an antigen-responsive population that is at least as long-lived as plasma cells.^45^ Memory B cells may play an important role in immune responses in tissues, including the lung mucosa and solid tumors,^46,47^ and can also re-seed the plasma cell compartment in response to repeat antigen exposure.^48,49^ The antigen-responsiveness of memory B cells could be exploited to support the amplification and persistence of engineered B cells, or to tune the levels of antibodies that are produced.

Many of these desirable features of an engineered B cell require that the cells maintain responsiveness to antigen, which can be achieved if the engineered antibodies maintain a BCR format. Genome editing allows reprograming of BCRs through site-specific insertion of custom antibody components at appropriate sites within the Ig locus. Most approaches take advantage of conserved sequences in the Eµ enhancer intron to insert cassettes comprising both a complete custom L chain and a custom V_H_ domain.^11–15^ The engineering of only IgH means that mismatch pairing is still possible with endogenous unedited L and H chains in the same cell that could create novel paratopes, and although endogenous Igκ can also be targeted for disruption,^11,14^ the structure of Igλ and its pseudogenes makes it much more challenging target to disrupt. In contrast, the HCAb editing approach described here offers a simpler path to B cell engineering, requiring only one DNA edit to insert a single polypeptide chain that can reprogram IgH for expression of a custom H chain antibody.

To validate the HCAb editing approach, we selected ARDs based on the J3 and A6 anti-HIV camelid V_H_H domains and inserted them into the intron downstream of the CH1 exon of IGHG1. This location ensures that the resulting H chain polypeptides exclude CH1 and will not interact with endogenous L chains. The engineered HCAbs retained the anti-HIV properties of the parental ARD, mirrored the physiology of a native IgG gene by regulating expression of both the membrane and secreted isoforms during *ex vivo* differentiation, and responded to antigen in a tonsil organoid model. Of interest, we also observed evidence that insertion of the J3 V_H_H at this locus supported SHM after extended culture in a B cell line, although the A6 V_H_H and a GFP control sequence did not. We note that our design of the J3 DNA sequence was codon wobbled from the published amino acid sequence^50^ using the closest human germline V_H_ (IGHV3-23) sequence as a template. In contrast, the A6 DNA sequence was unmodified from the natural antibody DNA sequence that had already undergone SHM evolution in its camelid host, which could explain the differing susceptibilities to hypermutation. The presence or absence of SHM could be uniquely advantageous in different settings. For instance, a variable pathogen such as HIV could be better controlled by HCAbs that retain the capacity to undergo SHM and subsequent affinity maturation, whereas a static target like a self-antigen may be better matched with an HCAb that does not hypermutate. Understanding the mechanisms that govern SHM at these insertion sites will require further research but may yield rules that allow tuning of sequences to either avoid or exploit this natural process of antibody evolution.

We further demonstrated that the simplicity of the HCAb design allows unprecedented customization of the engineered antibody. Antigen-recognition could be derived from a variety of different domains, both antibody and non-antibody, including V_H_Hs, an scFv domain, and a soluble receptor domain from CD4. The Fc region could additionally be programmed with additional substitutions by altering the site of insertion within the IgH constant regions. As an example, by moving the insertion site of J3 to be downstream of the CH2 domain in IGHG1, we successfully replaced both the ARD and the IgG1 Hinge and CH2 domains.

An ability to customize both the antigen-recognition and Fc components of HCAbs could significantly expand the potential applications of B cell engineering. Many diseases are currently treated with long-term monoclonal antibody infusions, in oncology, autoimmunity, neurological disorders, and beyond. Here, edited B cells could provide a long-lasting one-shot treatment, while the versatility of the HCAb editing platform could support many of the engineered enhancements that are included in these recombinant proteins.^1,2,20^ We hypothesize that the ARD component could accommodate a wide variety of different protein domains, including those from highly evolved antibodies, such as broadly neutralizing anti-HIV antibodies, or the antibodies recognizing self-antigens that are used to treat cancer or inflammatory disorders. The antigen-BCR response could even be driven by completely non-antibody components, such as receptor/ligand pairs. At the same time, the ability to substitute Fc components will allow the introduction of modifications to customize the effector functions of the antibody, such as GASDALIE mutations in CH2 that enhance ADCC activity.^51^ By moving the insertion site further downstream we can also substitute the CH3 domain (data not shown), which could allow the inclusion of mutations in this domain that extend antibody half-life^52^ or prevent interactions with endogenous H chains.^53,54^

While we have here focused on engineering in the IgG1 gene, this approach could also be applied to other Ig subclasses, creating HCAbs with specific desired properties, such as mucosal secretion conferred by IgA constant domains or the enhanced avidity of IgM pentamers. An advantage of editing within the constant region of a defined antibody subclass is that the HCAbs will not diversify through class switch recombination, but only express the selected engineered isotype. This could avoid the need for *ex vivo* or *in vivo* differentiation/immunization to drive isotype selection, while also generating a more consistent cell therapy product that will be stable over time and avoid potentially adverse effects of ongoing class switching.^55^ Conversely, while it remains possible class switch recombination events could loop out an edited HCAb, such cells would selected against by loss of amplification in response to the intended antigen-BCR interaction.

In conclusion, we have designed a simplified genome editing strategy to reprogram human B cells to express custom heavy chain antibodies, whose design accommodates customization of both the antigen-recognition component of the antibody and the Fc domain. It lends itself to a wide variety of possible designs, beyond conventional antibodies, as long as the final molecule can be expressed as a chimeric BCR-like molecule that can drive the naturally programmed responses of a B cell to its antigen.

## Methods

### DNA/RNA constructs and reagents

#### Antibody sequences and expression plasmids

J3,^24^ A6,^25^ CD4-mD1.22,^38^ and PGT121-scFv^39^ sequences were synthesized as gBlocks (IDT). The J3 protein sequence^24^ was first reverse translated (https://www.ebi.ac.uk/Tools/st/emboss_backtranseq/) with codon usage tables for *Homo Sapiens* and codon wobbled with silent mutations to match IGHV3-23*04, the closest human germline match identified by IgBLAST. PGT121 scFv was constructed with a standard (G_4_S)_3_ linker in the VH-VL orientation. J3, CD4-mD1.22, and PGT121-scFvs were paired with the leader from IGHV3-23D*01; A6 was paired with the leader from IGHV3-53*01.

The VH (3GIZ_H) and VL (3GIZ_L) sequences of the fully human anti-CD20 antibody Ofatumumab were obtained from NCBI. These VH and VL sequences, human IGKC, and the CH1 exon of human IGHG1 were codon optimized using the GeneOptimizer tool (Thermo Fisher) and synthesized as gBlocks (IDT). IGHV3-9 and IGKV3-11 leaders were placed upstream of VH and VL, respectively. The light chain comprised VL fused to IGKC. For expression of full-length Ofatumumab, the VL-IGKC cassette was inserted upstream of a P2A peptide, linked to VH-CH1, and followed by the remainder of the IGHG1 exons with native codon usage.

Expression plasmids for J3, A6, Ofatumumab-FL, and Ofatumumab-LC were generated in the pVAX backbone by Infusion cloning (Takara) of synthesized gBlocks (IDT). J3 and A6 sequences were additionally fused in frame to the Fc region of human IgG1 (including the upper hinge) to generate HCAb expression plasmids. Sequences of all expression plasmids and components are provided in Supplementary Table 2.

#### Guide RNAs

*Streptococcus pyogenes* Cas9 gRNAs with an NGG PAM targeting the human IGHG1 gene were designed using ChopChop (http://chopchop.cbu.uib.no) and CRISPOR (http://crispor.tefor.net) online tools. Sequences of gRNAs are reported in Supplementary Table 1. Synthetic single gRNAs with chemical modifications (2’-O-Methyl modifications at the first and last base and 3’ phosphorothioate bonds between the first 3 and last 2 bases) were synthesized by Synthego. Lyophilized gRNAs were resuspended at 100 µM in TE buffer (10 mM Tris, 1 mM EDTA, pH 8.0) and stored at -80°C.

#### Homology donors

Homology donors comprising sequences to be inserted and flanking homology arms were cloned into plasmid ITR-CMV-GFP (containing ITRs from AAV2; Cell Biolabs) or pVAX by Infusion cloning (Takara). The GFP expression cassette hPGK-eGFP-BGHpA was amplified by PCR from plasmid CCR5-PGK-GFP.^33^ Antibody components (J3, A6, CD4-mD1.22 or PGT121-scFv sequences, synthesized as gBlocks (IDT) as described above) were combined with the EEK promoter amplified from plasmid CCR5-EEK-GFP^33^ and the splice donor sequence from the IGHG1 CH1 exon.

All homology arms were symmetrical and of equal length, either 500 bp or 750 bp each. Homology arms to direct insertion downstream of the CH1 exon were amplified by PCR from human genomic DNA. Homology arms for insertion downstream of the CH2 exon were synthesized as gBlocks (IDT). Antibody cassettes to be inserted downstream of CH2 additionally contained sequences for the Hinge and CH2 exons of IgG1, which were codon optimized using GenSmart Codon Optimization tool (Genscript) to reduce homology with the endogenous sequences. The sequences of all homology donor plasmids and components are provided in Supplementary Table 3.

#### AAV vectors

AAV6-GFP homology donors were produced in-house as previously described, using triple transfection, tangential flow filtration, iodixanol gradient centrifugation, and ultrafiltration.^56^

The concentrated vectors were stored at -80°C. Vectors AAV6-sg05-750-EEK-J3-sd, AAV6-sg05- 750-EEK-PGT121-scFv-HL-sd, and AAV6-sg05-750-EEK-CD4-mD1.22-sd were produced by Vigene Biosciences. AAV6-CH2g1-750-EEK-J3-CH2*-sd vectors were produced by Vector Biolabs. AAV vectors are described in Supplementary Table 4.

All AAV vectors were titrated as previously described.^56^ Briefly, to remove residual plasmid DNA, vectors were treated with DNaseI (New England Biolabs) for 30 mins at 37°C, followed by inactivation at 75°C for 10 mins. AAV DNA was then extracted by treatment with proteinase K (Sigma-Aldrich) for 30 mins at 37°C, followed by inactivation at 95°C for 20 mins. The extracted DNA was stored at -20°C until titration. AAV vector genome (vg) titers were determined by TaqMan qPCR (Thermo Fisher) using ITR specific primers AAV ITR-Forward, AAV ITR-Reverse, and probe AAV ITR-Probe (5’-FAM, Tamra-3’) (Supplementary Table 5). To prepare the standard curve, serial dilutions of DNA extracted at the same time from a recombinant AAV2 Reference Standard Material (American Type Culture Collection; VR-1616) was used.^57^

### Antibody transfection and purification

#### HCAb production

HCAbs were generated by transient transfection of 293T cells (American Type Culture Collection, ATCC). Briefly, 4 x 10^6^ 293T cells were seeded into Poly-L-Lysine coated 10 cm dishes one day before transfection, in DMEM media supplemented with 10% FBS and 1% penicillin/streptomycin. The next day, 25 µg of expression plasmids pVAX-J3 or pVAX-A6 were introduced into cells using calcium phosphate transfection. After overnight incubation, cells were washed with PBS and media was replaced with serum-free UltraCULTURE media (Lonza) or phenol red-free DMEM with 10% FBS (Thermo-Fisher). Supernatants were harvested after 3 days, clarified by centrifugation for 15 mins at 2500 x *g* and filtered using a 0.45 µm filter. Antibodies were purified using Amicon Pro Affinity Concentration Kit Protein A with 10kDa Amicon Ultra-0.5 Device (Millipore-Sigma). Antibody concentrations were determined using an IgG ELISA, described below.

#### Co-transfections for cross-pairing assays

Antibody co-transfections to detect heavy and light chain pairings were performed as above, but using 2.5 x 10^5^ 293T cells seeded into each well of a 12-well plate one day before transfection and 1 µg of each plasmid (pVAX-J3, pVAX-Ofatumumab-FL or pVAX-Ofatumumab-LC), including for combinations. Supernatants were analyzed by ELISAs for gp120-V_H_H and gp120-Igκ, as described below.

### ELISAs

#### Total IgG ELISA

Protocols were adapted from previously described ELISA protocols.^58^ For the total IgG ELISA, 96 well high protein binding plates or strips (Corning 9018) were coated with polyclonal goat anti-human IgG antibody (Southern Biotech) in coating buffer (0.136% sodium carbonate, 0.735% sodium bicarbonate, pH 9.2) overnight at 4°C. Plates were washed 3 times with 350 µL wash buffer (PBS with 0.05% Tween-20) and blocked with 5% milk in wash buffer for 2-4 hours at 4°C. Standard curves were made with 1:2 serial dilutions of human IgG1, kappa from human myeloma (Millipore-Sigma) in blocking buffer in the range 2000-15.625 ng/mL. Samples were diluted in blocking buffer. Standards and samples were incubated overnight at 4°C, plates were washed 3 times with wash buffer, and 100 µL of anti-human IgG-Fc HRP (clone JDC-10, Southern Biotech), diluted 1:2000 in blocking buffer, was added to each well and incubated for 2 hours at 37°C. Plates were washed 3 times again, and detected with 100 µL of SigmaFast OPD (Millipore-Sigma) prepared according to the manufacturer’s instructions. Absorbance (450 nm) was measured using a Mithras LB 940 plate reader (Berthold Technologies).

#### gp120-V_H_H ELISA

The gp120-V_H_H ELISA was performed as described for total IgG, with the following modifications. Plates were coated with either 5 µg/mL HIV-1 JR-CSF gp120 (Immune Technology) in coating buffer for samples, or for the standard curve, with MonoRab™ Rabbit Anti-Camelid VHH Antibody (GenScript) diluted 1:250 in coating buffer. The standard curve was made using 1:2 serial dilutions in blocking buffer of the SARS-CoV-2 (COVID-19) Spike RBD Single Domain Antibody (ProSci, clone T4P3-B5), ranging from 2000-15.625 ng/mL. Samples were either subject to FabALACTICA digest or incubated with digestion buffer only (described below), and detected with MonoRab™ Rabbit Anti-Camelid VHH Cocktail [HRP] (Genscript), diluted 1:500 in blocking buffer.

#### gp120-Igκ ELISA

The gp120-Igκ ELISA was performed as described for total IgG, with the following modifications. Plates were coated with either 5 µg/mL JR-CSF gp120 (Immune Technology) in coating buffer for samples, or for the standard curve, directly coated with human IgG1, kappa from human myeloma (Millipore-Sigma), ranging from 2000-15.625 ng/mL by 1:2 serial dilutions in coating buffer. Samples were either subject to FabALACTICA digest or incubated with digestion buffer alone (described below), and detected with Mouse Anti-Human Kappa-HRP (SB81a) (Southern Biotech), diluted 1:4000 in blocking buffer.

#### gp120-IgG ELISA

The gp120-IgG ELISA was performed as described for total IgG, with the following modifications. Plates were coated with 5 µg/mL HIV-1 JR-CSF gp120 (Immune Technology) in coating buffer. A purified J3 HCAb standard (from 293T cell transfection, as described above) was quantified by total IgG ELISA and used to generate a standard curve ranging from 200- 1.5625 ng/mL by 1:2 serial dilutions in blocking buffer.

#### On-plate FabALACTICA digestion

2000 units of FabALACTICA enzyme (GENOVIS) were reconstituted in 50 µL water and diluted to 0.016 U/µL in digestion buffer (150 mM sodium phosphate buffer, pH 7.0 at 37 degrees). After overnight incubation of samples and washing, 50 µL of diluted enzyme was added to each well, or the digestion buffer alone to control wells. Digestion was carried out for 16 hours at 37°C. Plates were washed 8-10 times with 350 µL wash buffer per well, before continuing to detection steps.

### Cell culture and genome editing

#### Cell lines

K562, Raji, and Ramos 2G6 (CRL-1923) cells were obtained from ATCC. Cell lines were cultured in RPMI supplemented with 10% FBS and 1% penicillin/streptomycin. For gene editing, 2 x 10^5^ cells (K562) or 4 x 10^5^ cells (Raji and Ramos) were pelleted by centrifugation and washed with PBS. AAV6 transductions were performed in 10 µL of serum-free RPMI media for 1 hour at 37°C in a 96-well U-bottom plate prior to electroporation, as previously described.^33^ Cas9 RNPs were complexed by mixing 3000 ng TrueCut Cas9 protein v2 (Thermo Fisher) with 60 pmol of gRNA (Synthego) for 15 minutes at room temperature.

Cells were then washed with PBS and resuspended in 20 µL of SF (K562) or SG (Raji and Ramos) buffer (Lonza) and mixed with Cas9 RNPs. If appropriate, cells were also mixed with 1.4- 2 µg of plasmid homology donors or 100 pmol phosphorothioate-modified ssODNs as previously described.^59^ Electroporations were performed with the 4D-X Nucleofector (Lonza). K562 cells used pulse code FF-120, Raji cells used pulse code DS-104, and Ramos cells used pulse code CA-137.

J3-, A6-, or GFP-edited Raji and Ramos cells were expanded in culture. J3 and A6-edited cells were stained with anti-IgG-Fc antibody (BioLegend, clone M1310G05). Cells were sorted on a FACSAria II cell sorter based on membrane IgG expression (J3 and A6 editing) or GFP expression.

#### Primary B cells

Frozen human peripheral blood CD19^+^ B cells were purchased from StemCell Technologies and thawed per manufacturer’s instructions. IMDM (Corning) was supplemented with 10% FBS and 1% penicillin/streptomycin/amphotericin B. Immunocult-XF T cell expansion media (Stem Cell Technologies) was supplemented with 1% penicillin/streptomycin/amphotericin B and optionally with 10% FBS. Anti-RP105 (clone MHR73-1, BioLegend) was used at 2 µg/mL.^11,14^ Immunocult-ACF Human B Cell Expansion Supplement (BAC, Stem Cell Technologies) was used at 1:50 dilution per manufacturer’s instructions.

DP culture of B cells was as previously described.^33^ Briefly, cells were initially activated at 5 x 10^5^ cells/mL in B cell activation media: 5 µg/mL soluble CD40L (R&D Systems), 10 µg/mL anti-His tag antibody (clone AD1.1.10, R&D Systems), 50 ng/mL CpG ODN 2006 (Invivogen), 10 ng/mL IL-2 (R&D Systems), 50 ng/mL IL-10, and 10 ng/mL IL-15 (Peprotech). After 3 days of pre-activation, B cells were edited as described below and returned to B cell activation media for an additional 2 days. Then, cells were pelleted by centrifugation and media was replaced with plasmablast generation media: 10 ng/mL IL-2 (R&D Systems), 50 ng/mL IL-6 (Peprotech), 50 ng/mL IL-10, and 10 ng/mL IL-15. After a further 3 days of culture, cells were pelleted by centrifugation and media was replaced with plasma cell generation media: IMDM supplemented with 10% FBS, 50 ng/mL IL-6, 10 ng/mL IL-15, and 500 U/mL IFN-α (R&D Systems). Cells were cultured in this media for 3 additional days.

In all culture conditions, cells were edited after 3 days of pre-activation, as previously described.^33^ Briefly, cells were washed with PBS and resuspended at 5 x 10^7^ cells/mL in serum-free XF media. Ten µL of these cells were used for AAV6 transduction, at indicated MOIs, for 1 hour at 37°C in a 96-well U-bottom plate. Cas9 RNPs were complexed by mixing 15 µg TrueCut Cas9 protein v2 (Thermo Fisher) with 300 pmol of gRNA (Synthego) for 15 minutes at room temperature. Cells and AAV6 combinations were mixed with 90 µL BTXpress Electroporation Buffer and RNPs, transferred to Electroporation Cuvettes Plus (2 mm gap, BTX), and electroporated using a BTX ECM 830 at 250 V for 5 ms. After electroporation, cells were diluted to 5 x 10^5^ cells/mL in original culture media.

#### Tonsil organoids and immunization

Tonsil tissue was received from the Department of Otolaryngology-Head & Neck Surgery, Keck School of Medicine of the University of Southern California, as anonymous waste samples, approved by USC’s Institutional Review Board (protocol HS-17-00023-AM001). Tonsil mononuclear cells (TMNCs) were harvested as previously described.^37^ Briefly, tonsils were dissected into small pieces and mashed into a single cell suspension using 70 µm cell strainers. TMNCs were purified via Ficoll density gradient separation (700 x *g*, 20 min, no brake). B cells were purified from TMNCs with the EasySep Human CD19 Positive Selection Kit II (Stem Cell Technologies) per the manufacturer’s instructions. TMNCs or purified B cells were frozen in Cryostor (Stem Cell Technologies) and stored in liquid nitrogen until use.

Tonsil B cells were cultured and edited using the same protocol as for PBMC-derived B cells, with BAC and serum-free XF media. Two to 4 days post-editing of the B cells, the matched TMNC aliquots were thawed, enumerated, and combined with populations of edited or unedited tonsil B cells in 19:1 ratio of TMNCs to B cells to create the tonsil organoid. Final cell numbers were 6 x 10^7^ cells/ml for large cultures (24-well size transwell plate, Millipore-Sigma) or 2 x 10^7^ cells/ml for small cultures (96-well size transwell plate, Millipore-Sigma). 100 µL of cell suspension was plated into permeable membranes in transwell plates, and tonsil culture media was added to the basolateral chamber (1 mL for 24-well plates, 300 µL for 96-well plates). Tonsil culture media comprised RPMI with GlutaMAX (Gibco) supplemented with 10% FBS, 1% penicillin/streptomycin, 1% MEM Non-Essential Amino Acids Solution (Gibco), 1 mM Sodium Pyruvate (Gibco), 100 µg/mL Normocin (InvivoGen), 1% Insulin-Transferrin-Selenium (ITS-G, Gibco), and 200 ng/mL recombinant human B cell-activating factor (BAFF, BioLegend).

For immunization, 1 µg (24 wells) or 0.33 µg (96 wells) recombinant HIV-1 JR-CSF gp120 (Immune Technology) or control antigen PE (Thermo Fisher) were mixed with adjuvants per manufacturer’s protocols and added to cells immediately after plating. Alhydrogel 2%, Adju-Phos, AddaVax, and Quil-A adjuvants were all from Invivogen. Culture medium was changed and supernatants harvested every 2 to 3 days until cell harvest.

### Assays for genome editing

#### Cell viability and counting

Cells were mixed with Guava ViaCount Reagent (Luminex) at a 1:20 ratio. Cell concentrations and viability were determined using the ViaCount assay using a Guava easyCyte machine (Millipore-Sigma).

#### Flow cytometry

All cell staining was performed in PBS. For cells stained with gp120, 1 µg of His-tagged HIV-1 JR-CSF gp120 (Immune Technology) was added to cells for 15 min at room temperature. Cells were washed with PBS, then stained with fluorescent anti-His antibody (Miltenyi Biotec), along with other antibodies according to panels described in Supplementary Table 6. Data was acquired using a FACSCanto II (BD Biosciences), FACSAria II (BD Biosciences), Attune NxT Flow Cytometer (Thermo Fisher), or Guava easyCyte (Millipore-Sigma). Data were analyzed using FlowJo software (FlowJo, LLC).

#### HIV neutralization assay

Stocks of HIV-1 strains JR-CSF and NL4-3 were produced by plasmid transfection of 293T cells, as previously described.^60^ HIV neutralization activity was measured for serial dilutions of the supernatants from J3 or A6-edited B cell lines using the TZM-bl assay, as previously described.^61^ Luminescence was detected with the britelite plus kit (Perkin-Elmer) and measured with a Mithras LB 940 plate reader (Berthold Technologies). Control J3 and A6 HCAbs were generated from transfected 293T cells. Input amounts of each virus were titrated to give roughly 10-20 times higher luminescence than background in control wells receiving no virus or antibodies.

#### ICE assay

Cells were pelleted and genomic DNA was extracted using the DNeasy Blood & Tissue Kit (Qiagen). Roughly 200 ng of genomic DNA was subjected to PCR using primers specific for IGHG1, IGHG2, IGHG3, IGHG4, or IGHGP with AmpliTaq Gold 360 Master Mix (Thermo Fisher). Primer sequences are provided in Supplementary Table 5. The PCR reaction was run using touchdown PCR^62^ on a C1000 Touch Thermal Cycler with the following conditions: 95°C 10 min, 15 cycles (95°C 30 s, 70°C 30 sec [-1°C/cycle], 72°C 1 min), 20 cycles (95°C 30 s, 55°C 1 min, 72°C 1 min), 72°C 7 min, and 4°C forever. For IGHG3, the extension time was 1 min 40 sec due to a longer PCR product. All PCR products were Sanger sequenced using the primer IGHG-UniSeq-F2 by Genewiz, and the output was analyzed using the ICE online tool (https://ice.synthego.com).^63^

#### In-out PCR

Cells were pelleted and genomic DNA was extracted using the DNeasy Blood & Tissue Kit (Qiagen). Roughly 50-250 ng of genomic DNA was subject to PCR using primers IGHG1-HA-Fwd2 and IO-EEK-In-6 (Supplementary Table 5). PCR products were visualized on a 1% agarose gel with GelRed Nucleic Acid Stain (Biotium). PCR products were purified using Nucleospin Gel and PCR Clean-up kit (Takara), and Sanger sequenced using primer IGHG1-F1 (Supplementary Table 5) by Genewiz. Chromatograms were plotted in R using the sangerseqR package.^64^

#### RT-PCR

RNA was isolated from cell pellets using Nucleospin RNA Plus XS kit (Takara), and first strand cDNA synthesis was performed using Superscript IV VILO Master Mix with ezDNAse (Thermo Fisher) per manufacturer’s instructions. PCR was performed using primers J3-RT-F2 and IGHG1-Sec-RT-R2 (secreted H chain isoforms) or IGHG1-MemEx-RT-R1 (membrane anchored H chain isoforms) (Supplementary Table 5), using 5 µL of cDNA. Resulting DNA was visualized on a 1% agarose gel with GelRed Nucleic Acid Stain (Biotium).

#### Droplet digital (dd)PCR

Genomic DNA from edited cells was extracted using the DNeasy Blood & Tissue Kit (Qiagen) and the percent edited alleles measured for an in-out PCR product, amplified with primers IGHG1-HA-Fwd2, IO-EEK-In-6, and probe sg05-P1 (5’ FAM; ZEN/3’-IBFQ double quenched) (Supplementary Table 5). A human RPP30 copy number assay labeled with HEX (Bio-Rad, dHsaCP1000485) was used to measure total allelic copy numbers. Droplets were prepared using ddPCR Supermix for Probes (No dUTP) and a QX200 Droplet Generator (Bio-Rad). The PCR reaction was run on a C1000 Touch Thermal Cycler with the following conditions: 95° C 10 mins (ramp 2°C/sec), 40 cycles [94°C 1 min (ramp 1°C/sec), 55°C 30 sec (ramp 2°C/sec), 72°C 2 min (ramp 1°C/sec)], 98°C 10 min (ramp 2°C/sec), and 4°C forever (ramp 2°C/sec). Droplets were read on a QX200 Droplet Reader and the copy number variation data was analyzed with QuantaSoft analysis software (Bio-Rad).

### Next-generation sequencing

#### sg05 activity in primary B cells

Genomic DNA from primary B cells treated with sg05 RNPs was isolated using the DNeasy Blood & Tissue Kit (Qiagen) and amplified for sequencing by nested PCR using Platinum SuperFi II Master Mix (Thermo Fisher). The DNA was initially amplified with primer pairs specific for each gene (IGHG1, IGHG2, IGHG3, IGHG4, or IGHGP; see Supplementary Table 5) for 15 cycles to enrich for the gene to be amplified. Secondary PCR using primers AEZ-IGHG1-Fwd and AEZ-IGHG1-Rev for 35 cycles was used to amplify DNA and add partial Illumina adapters for sequencing, including in-line barcodes to minimize index hopping (Supplementary Table 5).^65^

Next generation sequencing (2 x 250 bp paired end) was performed using the Amplicon-EZ service (Genewiz). FASTQ files were aligned and filtered using PANDAseq,^66^ imported into RStudio and analyzed using the ShortRead^67^ and Biostrings^68^ packages. Mutated reads were identified based on insertions, deletions, >2 bp changes, or combinations thereof. To calculate mutations by position around the sg05 cut site, filtered reads were aligned pairwise with the reference sequence, and a position weight matrix was calculated based on these alignments. Results were exported using the openxlsx package.^69^

#### Somatic hypermutation in B cell lines

Isolated genomic DNA from J3, A6 or GFP-edited Raji cell populations was amplified by an initial in-out PCR. J3 and A6 used primers EEK-End-Fwd and IGHG1-HA-Rev1 (Supplementary Table 5) for 15 cycles, to amplify sequences inserted specifically at IGHG1. Secondary PCR using primers AEZ-J3-Fwd or AEZ-A6-Fwd and AEZ-VHHsd-Rev (Supplementary Table 5) for 35 cycles was used to amplify DNA and add partial Illumina adapters for sequencing. GFP was amplified initially with primers PGK-Seq-Fwd and IGHG1-HA-Rev1, followed by amplification of the first 400 bp of GFP and addition of partial Illumina adapters using primers AEZ-GFP-Fwd and AEZ-GFP-400-Rev. Sequencing and import were performed as above. AID hotspots were identified as WRCH motifs in each sequence.^70^ Sequence logo plots were generated using the ggseqlogo package.^71^

### Statistics

Data are expressed as mean ± SEM. Inter-group differences were assessed as appropriate and indicated in figure legends. Statistical tests were calculated in Prism 9.5.1 (Graphpad) and included 2-tailed unpaired Student’s t-test, 2-tailed paired Student’s t-test, 1- sample t-test, 1-way ANOVA, 2-way ANOVA, Pearson’s correlation, 3 parameter nonlinear regression, and asymmetric 5-paramater nonlinear regression. Hypothesis tests were 2-sided, and the threshold of significance was set to 0.05.

## Acknowledgements

We thank George N. Llewellyn and Catherine Diadhiou for critical discussions and assistance reviewing the manuscript. This work was supported by National Institutes of Health (NIH) grants HL156274, AI164561, AI164556 and MH130178 to P.M.C. G.L.R. was supported by a Career Development Award from the American Society of Gene & Cell Therapy (ASGCT). H.- Y.C. was supported by a Taiwan USC scholarship. Tonsil material was provided by the Norris Comprehensive Cancer Center Translational Pathology Core, supported in part by NIH grant CA014089 from the National Cancer Institute (NCI). The content is solely the responsibility of the authors and does not necessarily represent the official views of ASGCT, NCI or NIH.

## Author contributions

G.L.R., C.H., K.S., A.M., H.M., and H.-Y.C developed editing reagents and performed cell editing experiments. G.L.R., C.H., A.M., X.H., K.S., and C-H.C. established assays and performed antibody and sample analyses. G.L.R. performed NGS experiments, analyzed data, and visualized results. E.J.K. supervised collection of tonsil tissue. G.L.R. and P.M.C. conceived the study, designed experiments, interpreted data, and wrote the manuscript with input from other authors. P.M.C. supervised the study.

---

**Extended Data Figure 1.**
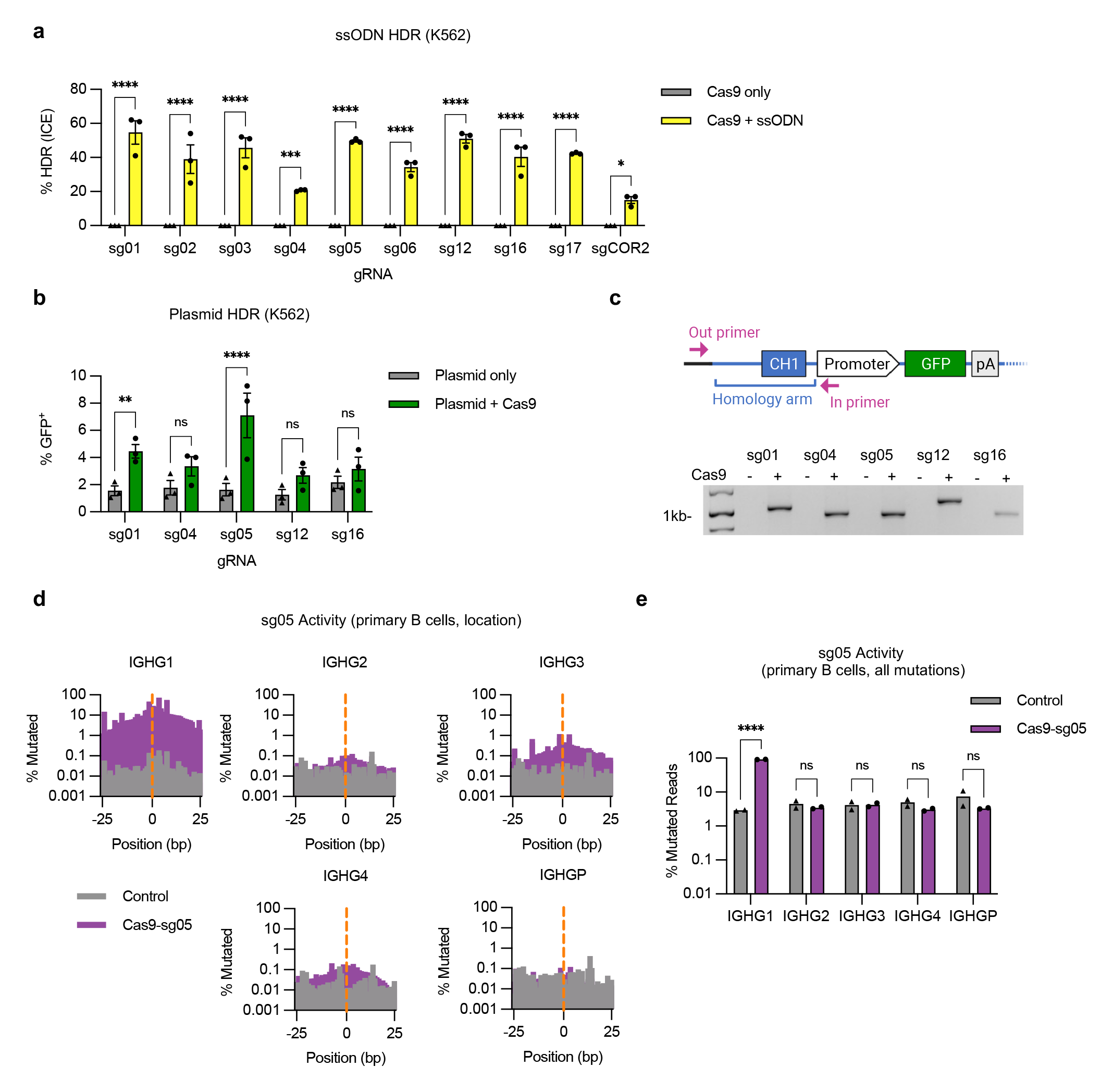
Extended analyses of genome editing at the constant region of the IgH locus. (**a**) K562 cells were electroporated with Cas9 RNPs containing indicated gRNAs and matched ssODN homology donors to insert an XhoI restriction site (*n* = 3). HDR editing was measured by Sanger sequencing and ICE analysis. (**b**) K562 cells were electroporated with Cas9 RNPs for indicated gRNAs and a matched plasmid homology donor containing a GFP expression cassette (*n* = 3). HDR editing was measured by flow cytometry for GFP expression after 3 weeks. (**c**) Site-specific insertion of GFP expression cassettes in AAV6-edited K562 cells was confirmed by in-out PCR for each tested gRNA. Uncropped gel is available in Supplementary Fig. 3a. (**d**) On-and off-target activity of sg05 was measured at indicated IGHG genes in primary human B cells, 5 days after editing, by targeted amplicon deep sequencing. Aggregate mutations at each base in a 50bp window surrounding the sg05 cut site (0; orange dotted line) are shown for each gene. (**e**) Percentage mutated reads at each IGHG gene calculated as for all changes (≥ 1 bp changed), which gives a higher background than when a cutoff of ≥ 2 bp is selected, as shown in Fig. 1d (*n* = 2). Error bars show mean ± SEM. Statistics were calculated by 2-way ANOVA. * *p* < 0.05, *** *p* < 0.001, **** *p* < 0.0001, ns = not significant.

**Extended Data Figure 2.**
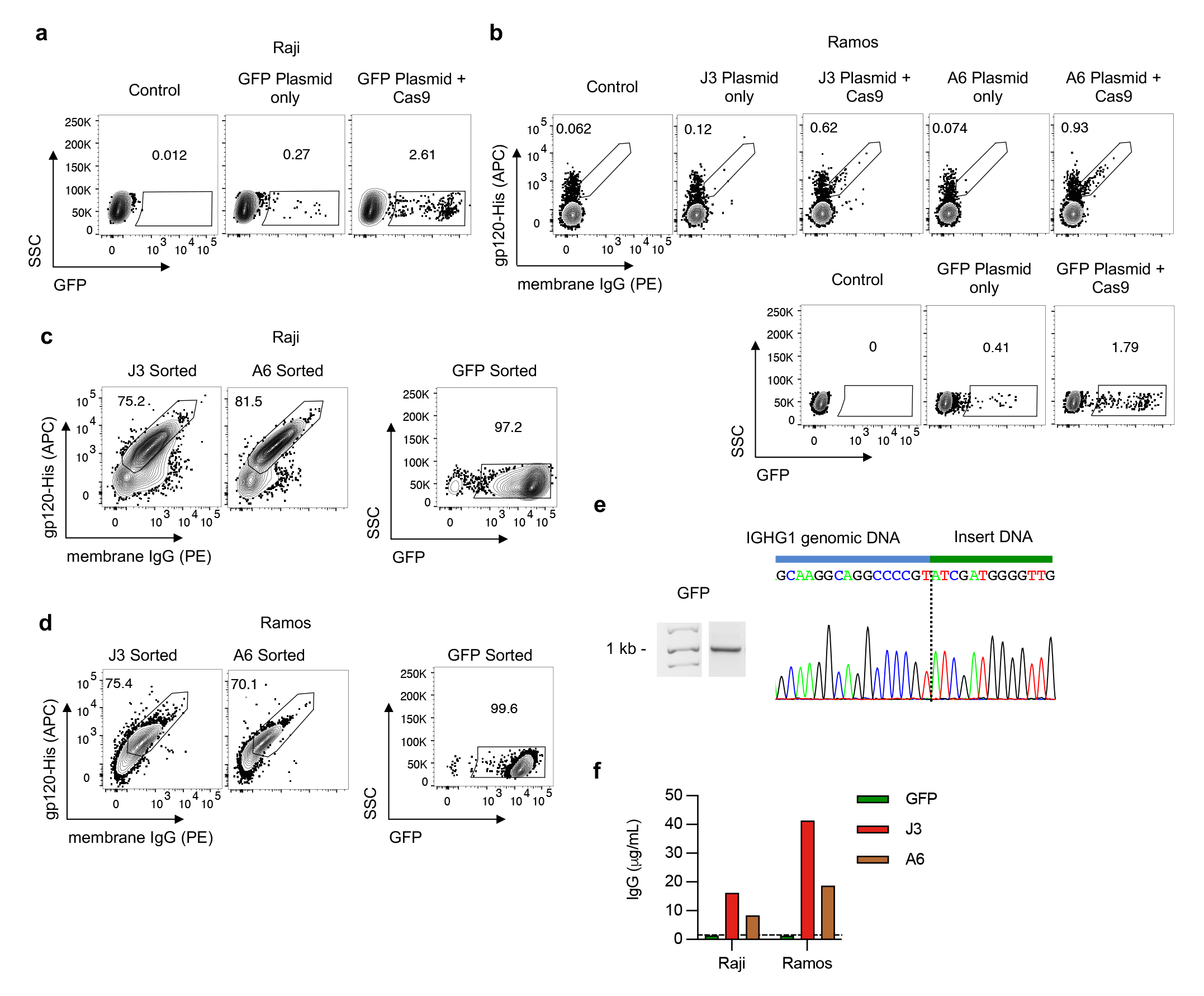
Extended analyses of engineered B cell lines expressing anti-HIV HCAbs. (**a**) Raji B cells were edited with sg05 Cas9 RNPs plus a plasmid homology donor to insert a GFP expression cassette and editing rates were measured by flow cytometry. (**b**) Ramos B cells were edited with sg05 Cas9 RNPs and plasmid homology donors for J3, A6 or a control GFP expression cassette. Editing rates were determined by flow cytometry. (**c-d**) Raji (**c**) and Ramos (**d**) cells edited by sg05 Cas9 RNPs and J3, A6, or GFP plasmid homology donors, post sorting by FACS. (**e**) GFP-edited Raji cells were sorted by FACS for GFP expression, and the enriched population was subjected to in-out PCR and Sanger sequencing of PCR bands to confirm precise insertion. The dotted line indicates the predicted sg05 cut site. Uncropped gel is available in Supplementary Fig. 3b. (**f**) Secretion of HCAbs was detected by total IgG ELISA from J3 or A6-edited Raji and Ramos cells, but not from GFP-edited control cells.

**Extended Data Figure 3.**
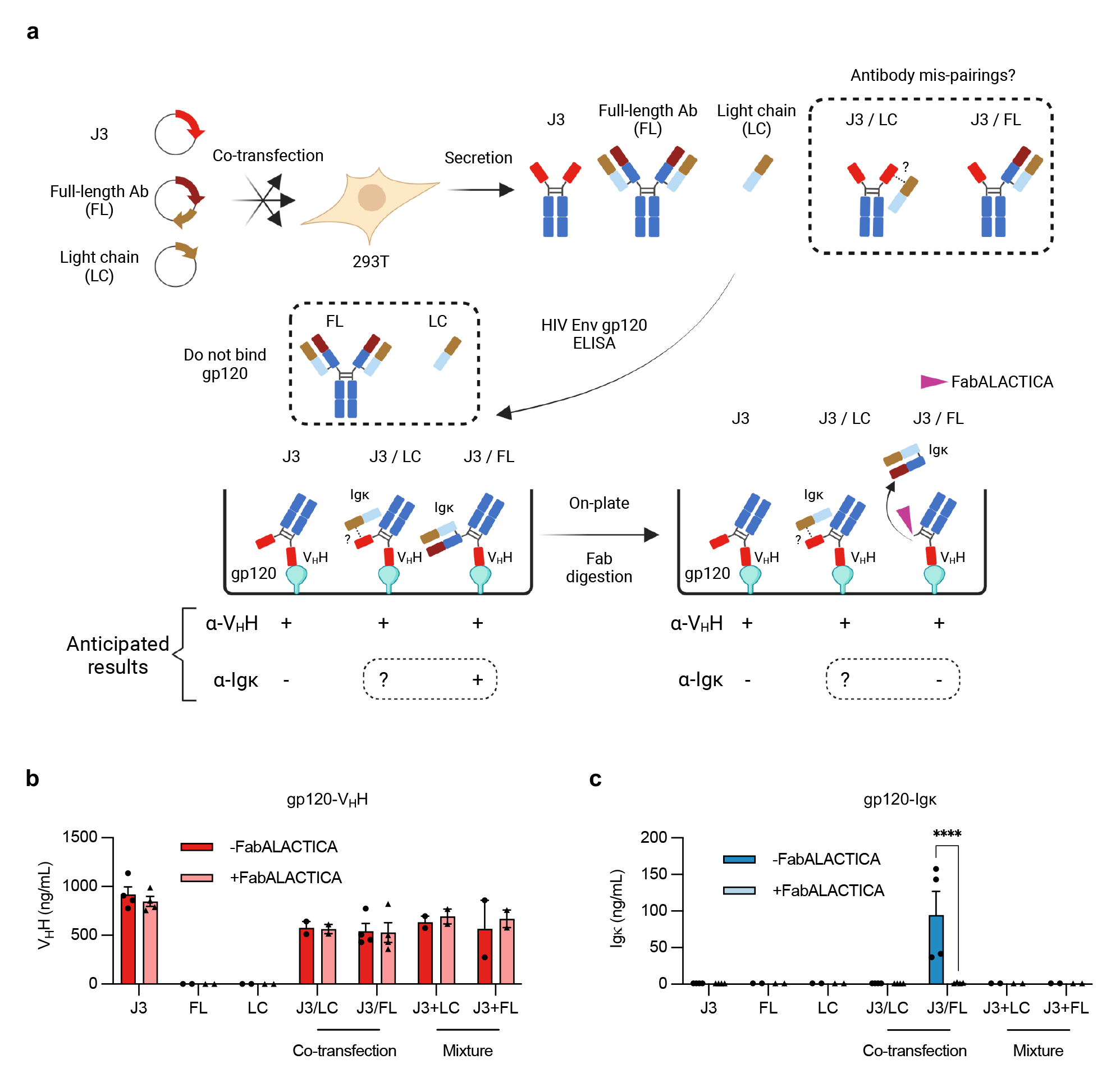
Lack of pairing between HCAb and co-expressed light chain. (**a**) Assay schematic. 293T cells were transfected with expression plasmids for the J3 V_H_H HCAb, a full-length (FL) conventional human IgG1 derived from Ofatumumab, or its Igκ L chain only (LC), or combinations as indicated. Supernatants, including mixed supernatants as controls, were evaluated by ELISAs based on HIV gp120 binding by the J3 V_H_H and detection with an anti-V_H_H antibody (control), or anti-Igκ L chain antibodies to detect pairings between the J3 HCAb and either the FL or LC components. FabALACTICA digestion is expected to cleave FL antibodies into Fab and Fc fragments and may also cleave HCAbs. (**b-c**) Results of gp120-V_H_H (**b**) and gp120-Igκ (**c**) ELISAs, with or without FabALACTICA digestion *(n* = 2-4). Cross-pairing was only observed after co-transfection of J3 HCAb and FL antibody, consistent with H chain interactions that were released by FabALACTICA digestion. In contrast, the L chain alone did not pair with the J3 HCAb. Error bars show mean ± SEM. Statistics were performed using 2-way ANOVA. **** *p* < 0.0001.

**Extended Data Figure 4.**
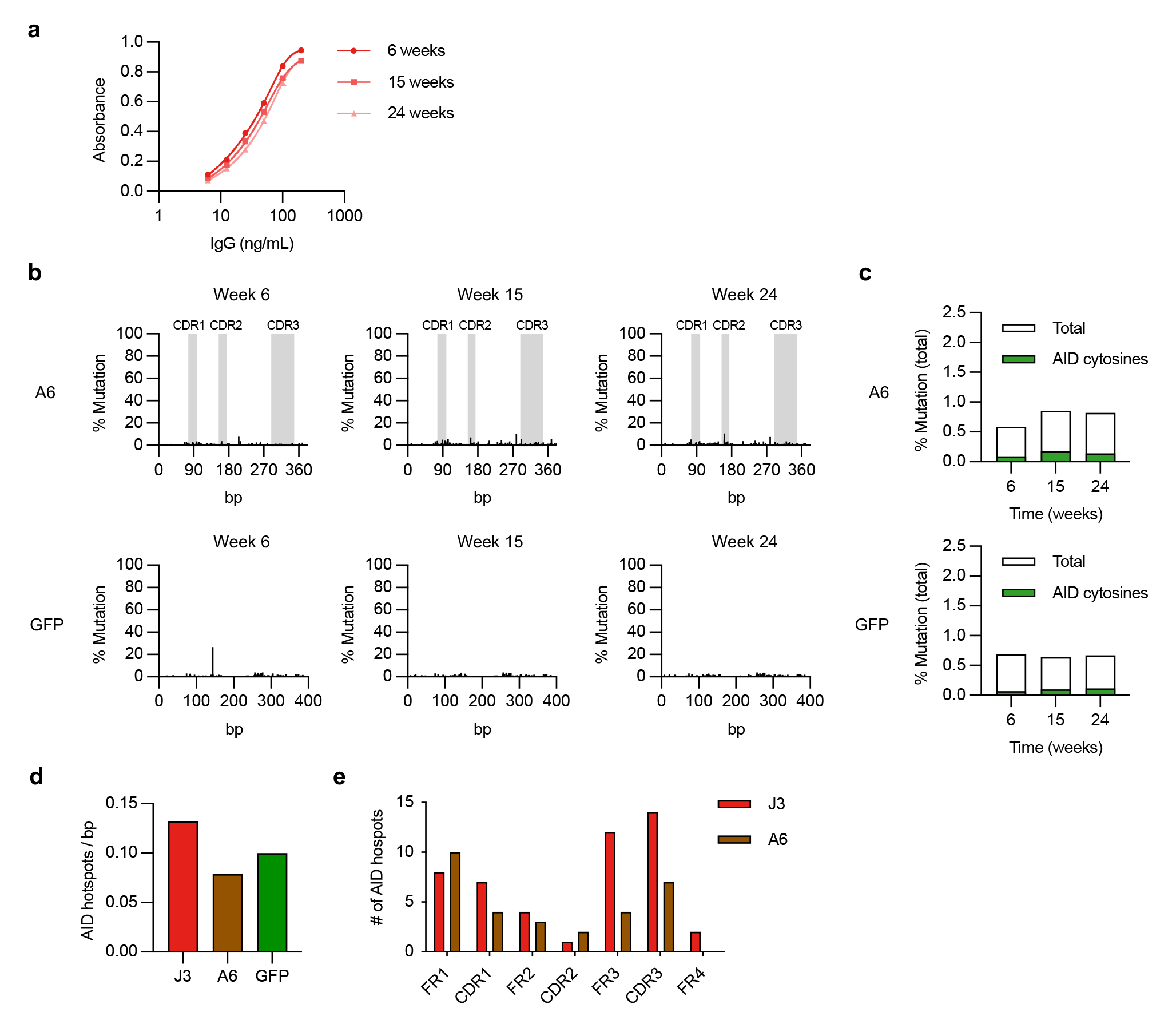
Lack of somatic hypermutation in inserted A6 V_H_H and GFP sequences. (**a**) Absorbance curve for J3 avidity over time from J3-edited Raji cells in extended culture (*n* = 3). IgG produced by the cells at indicated time points was measured by gp120-IgG ELISA. (**b**) Changes at A6 or GFP sequence in edited Raji cells over time, measured by deep sequencing. Percentage mutation at each position is the frequency of reads that did not match the wild-type sequence. CDR regions in A6 are indicated in grey. (**c**) Total mutations in A6 or GFP sequences at each timepoint were summed and divided by the total sequence length to determine a total % mutations. Shown in green are mutations associated with AID hotspot motif cytosines (WR<u>C</u>H). (**d**) The density of AID hotspots (number of WRCH hotspots / number of base pairs in the sequence) for each indicated sequence. (**e**) The distribution of AID hotspots across framework regions (FR) and complementarity-determining regions (CDR) for J3 and A6. Error bars show mean ± SEM.

**Extended Data Figure 5.**
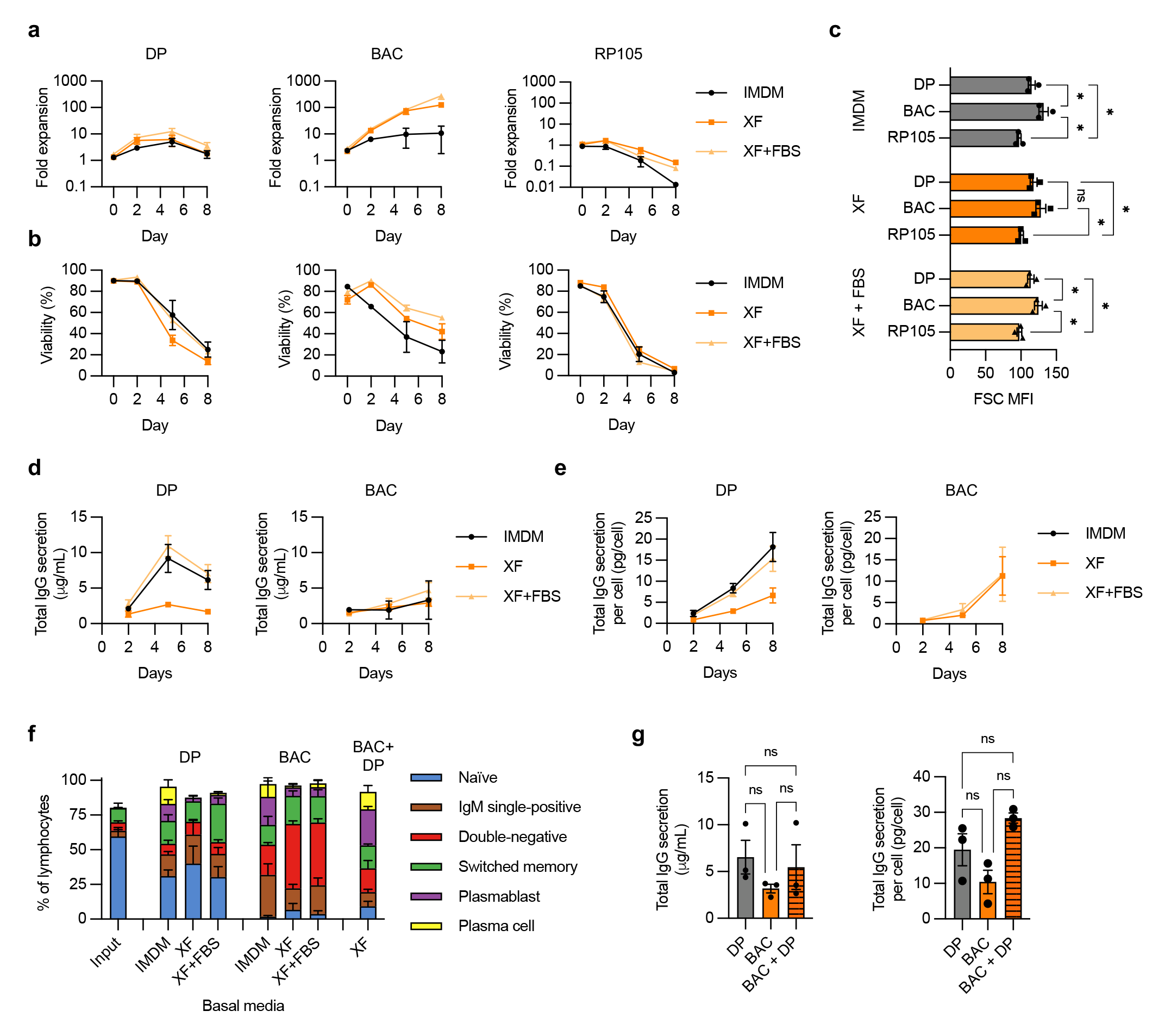
Comparison of culture conditions for primary human B cells. Primary human B cells from *n* = 3-4 independent experiments were cultured with the indicated stimulation conditions (DP, BAC, RP105) and basal media (IMDM, XF, XF plus FBS) as shown in Supplementary Fig. 4. Cultures were pre-activated for 3 days before measurements were started on day 0. (**a**) Fold-expansion of cells over time. (**b**) Viability of cells over time. (**c**) Cell size, measured by flow cytometry as forward scatter (FSC) median fluorescence intensity (MFI) on day 0. (**d**) Total IgG secreted into supernatants was measured over time by ELISA. (**e**) Total IgG amounts normalized for viable cell counts. (**f**) B cell phenotypes were assessed at day 8 in indicated cultures by flow cytometry, as shown in Supplementary Fig. 5. For BAC+DP, the cells were started in BAC and treated for a total of 5 days, then switched to the DP protocol for the remaining 6 days, as shown in Supplementary Fig. 4. (**g**) Comparison of total IgG secretion and IgG/cell at day 8 for cells treated with indicated stimulation protocols. DP was in IMDM, BAC and BAC + DP were in XF without FBS. Error bars show mean ± SEM. Statistics in panels (c,g) were calculated by one-way ANOVA. * *p* < 0.05, ns = not significant.

**Extended Figure 6.**
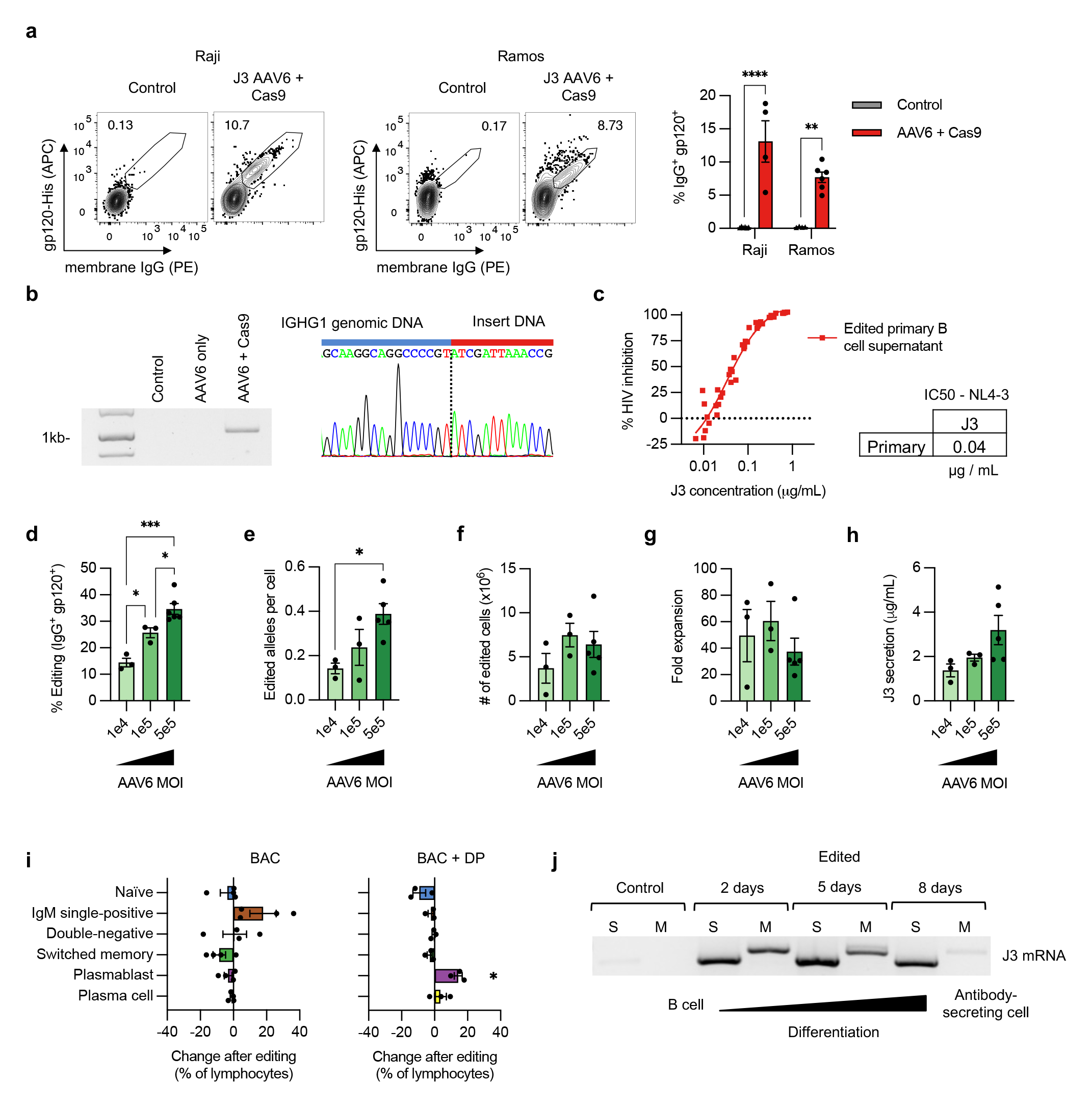
Extended analyses of engineering with AAV6 homology donors. (**a**) Editing rates achieved using sg05 Cas9 RNPs and AAV6-J3 homology donors on Raji and Ramos B cell lines. Representative plots are shown, together with summary data for *n* = 4-6 replicates. (**b**) Site-specific insertion of the J3 cassette in primary B cells treated with sg05 RNPs and AAV6-J3 donor was confirmed by in-out PCR followed by Sanger sequencing of the band. The dotted line shows the presumed sg05 cleavage site. Uncropped gel image is provided in Supplementary Fig. 3c. (**c**) Neutralization of HIV-1 NL4-3, measured by TZM-bl assay, by supernatants of AAV6- J3-edited primary B cells *(n* = 6). (**d-h**) Primary human B cells from *n* = 3-5 donors were edited with sg05 Cas9 RNPs and AAV6-J3 at the indicated MOIs, with BAC activation in XF media. Data from the highest MOI is reproduced here from Fig. 4, for comparison. (**d**) Editing rates, measured at day 8 by flow cytometry for surface J3- BCR. (**e**) Editing quantified by in-out ddPCR at day 8 and normalized per cell against a control reaction. (**f**) The yield of edited cells at day 8 was calculated from 5 x 10^5^ starting B cells. (**g**) Fold expansion of total cells at day 8. (**h**) J3 HCAb secretion, measured by gp120-IgG ELISA at day 8. (**i**) B cell phenotypes of edited cells in BAC (left) or with differentiation (BAC + DP, right) were measured by flow cytometry. Unedited matched control cells were also quantified and subtracted from the frequencies in the edited cells, to highlight any changes in differentiation after editing. (**j**) Edited primary B cells were differentiated in DP plus IMDM media and RNA extracted at indicated time points. RT-PCR was performed using a J3-specific forward primer and IgG reverse primers specific for the secreted (S) or membrane (M) isoforms. Uncropped gel image is provided in Supplementary Fig. 3d. Error bars show mean ± SEM. Statistics were calculated by 2-way ANVOA (a), 1-way ANOVA (d-h), or 1-sample t-test (i). * *p* < 0.05, ** *p* < 0.01, *** *p* < 0.001, **** *p* < 0.0001.

## Notes

### Competing Interest Statement

The authors have declared no competing interest.

