## Supplementary Figures 1-8 for "Reprogramming human B cells with custom heavy chain antibodies"

**Supplementary Figure 1**

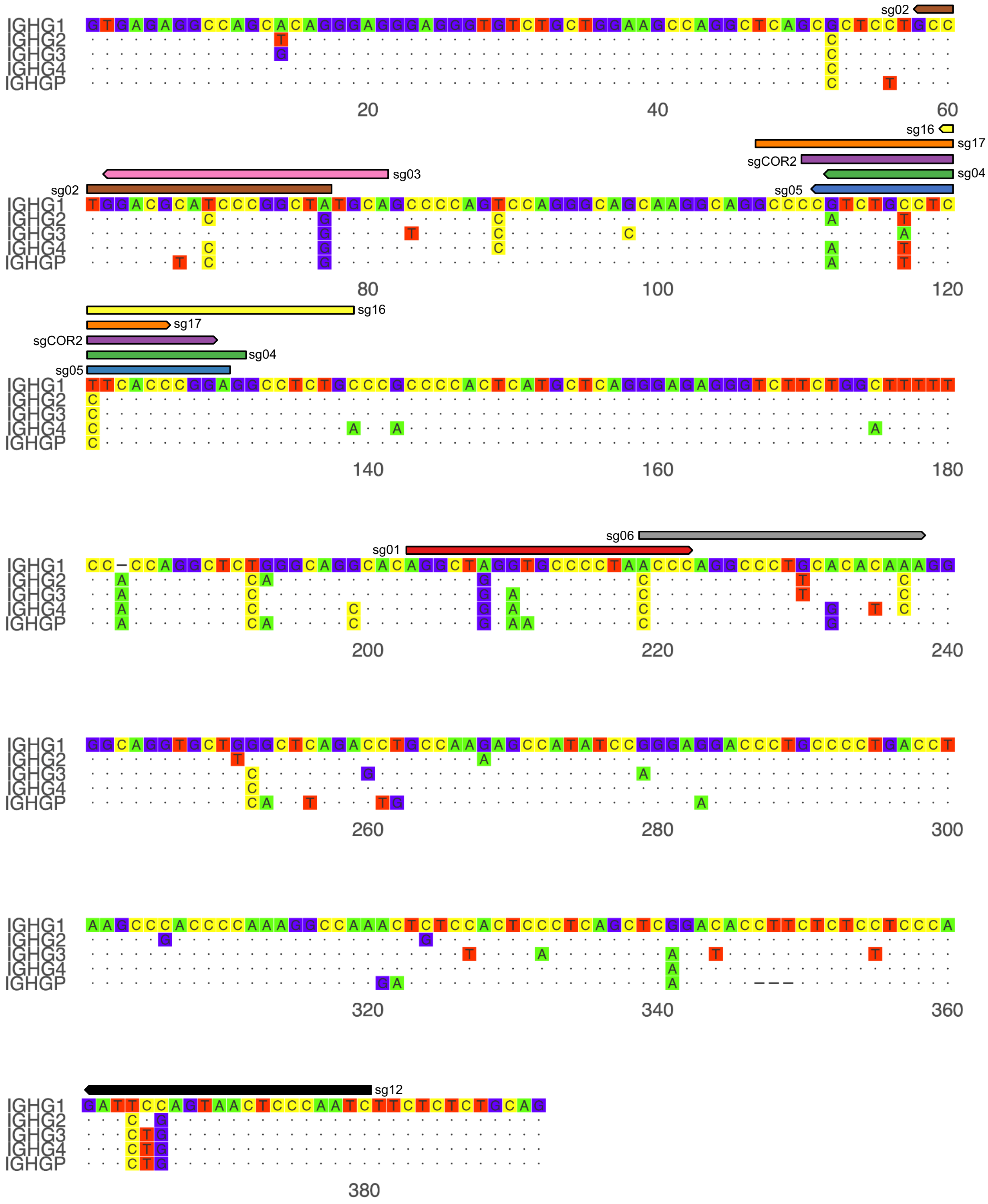


**Supplementary Figure 1. Alignment of CH1 intron sequences of IgG constant regions.** The sequence of the intron downstream of CH1 is aligned for IgG1 (IGHG1) and other IgG genes. Designed spCas9 guide RNA sequences are shown above the sequence, with the arrow direction indicating the sequence (right-pointing arrows have the 5'-3' sequence shown, left-pointing arrows are the complementary sequence).

### Supplementary Figure 2

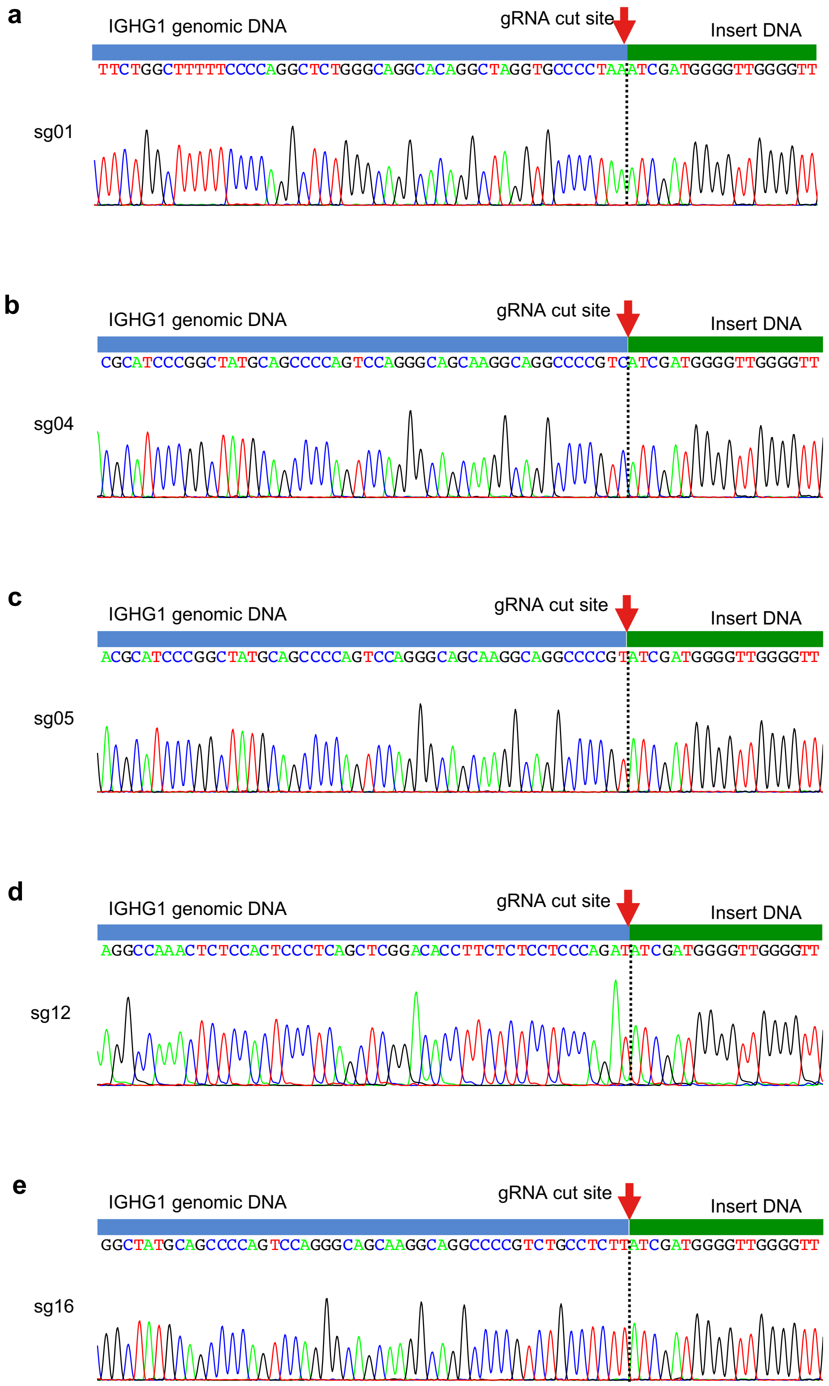


**Supplementary Figure 2. Sequencing of in-out PCR products from HDR-edited K562 cells.** In-out PCR products from the indicated gRNAs in Extended Data Fig. 1c were subjected to Sanger sequencing. Each sequence trace is annotated to indicate the IGHG1 genomic DNA sequence, the predicted gRNA cut site, and the start of the inserted sequence.

### Supplementary Figure 3

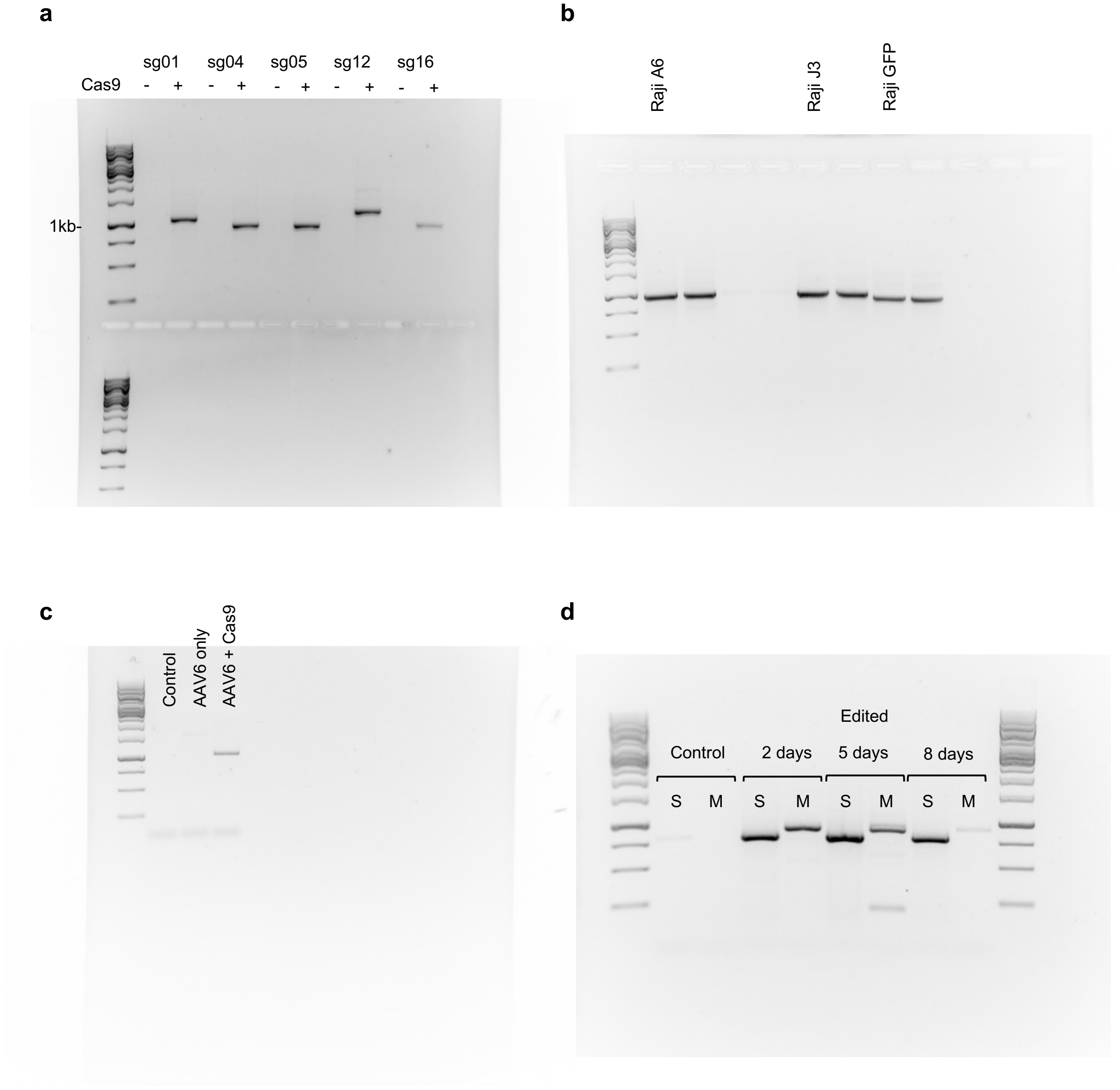


**Supplementary Figure 3. Uncropped gel images.** (**a**) For Extended Data Fig. 1c. (**b**) For Figure 2b and Extended Data Fig. 2b. (**c**) For Extended Data Fig. 6b. (**d**) For Extended Data Fig. 6j. Any unlabeled lanes contain irrelevant samples from other experiments.

### Supplementary Figure 4

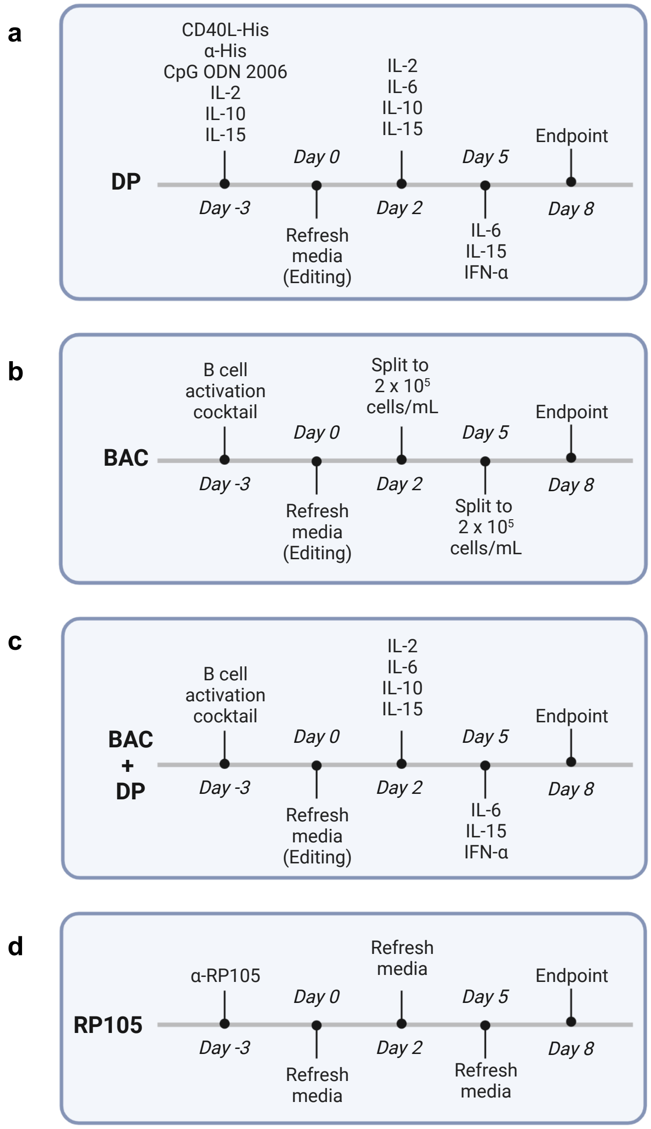


**Supplementary Figure 4. Culture protocol timelines.** Timelines are shown for culture with (**a**) the differentiation protocol (DP), (**b**) B cell activation cocktail (BAC), (**c**) BAC plus subsequent differentiation (BAC + DP), or (**d**) anti-RP105 activation (RP105). If appropriate, editing was performed at day 0, after 3 days of pre-activation. Where "refresh media" is indicated, media was completely removed from cells after centrifugation, and cells were resuspended in an equal volume of fresh media containing the same media/supplements as indicated previously. For BAC, cells were also split as indicated, with complete media replacement.

### Supplementary Figure 5

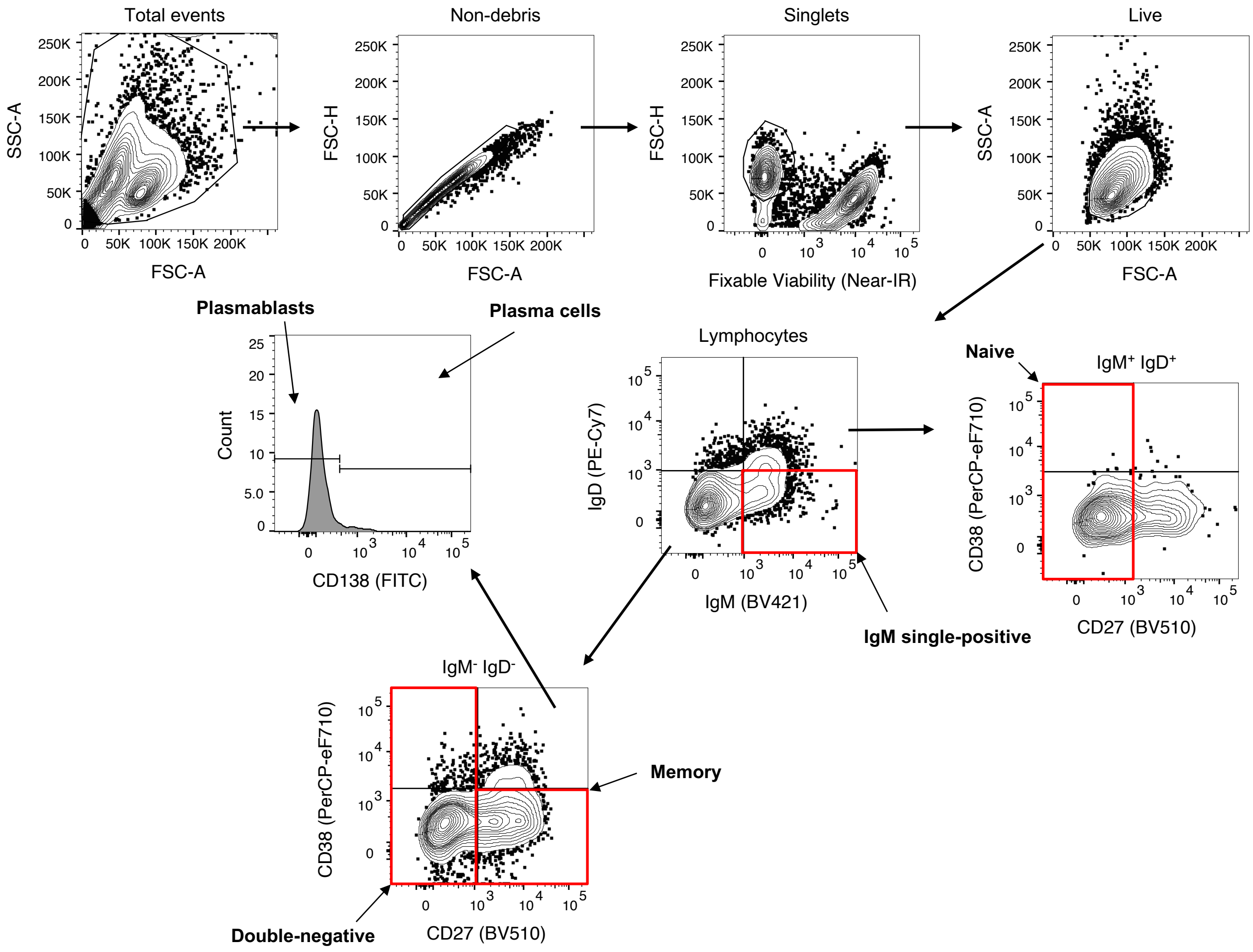


**Supplementary Figure 5. Representative gating for analysis of B cell differentiation.** Total lymphocytes were identified by excluding debris, non-singlets, dead cells, and on the basis of size. Naïve B cells are $IgM^{+}$ $IgD^{+}$ $CD27^{-}$ $CD38^{-/+}$. Double-negative B cells are $IgM^{-}$ $IgD^{-}$ $CD27^{-}$ $CD38^{-/+}$. Memory B cells are $IgM^{-}$ $IgD^{-}$ $CD27^{+}$ $CD38^{-}$. Plasmablasts are $IgM^{-}$ $IgD^{-}$ $CD27^{+}$ $CD38^{+}$ $CD138^{-}$. Plasma cells are $IgM^{-}$ $IgD^{-}$ $CD27^{+}$ $CD38^{+}$ $CD138^{+}$.

### Supplementary Figure 6

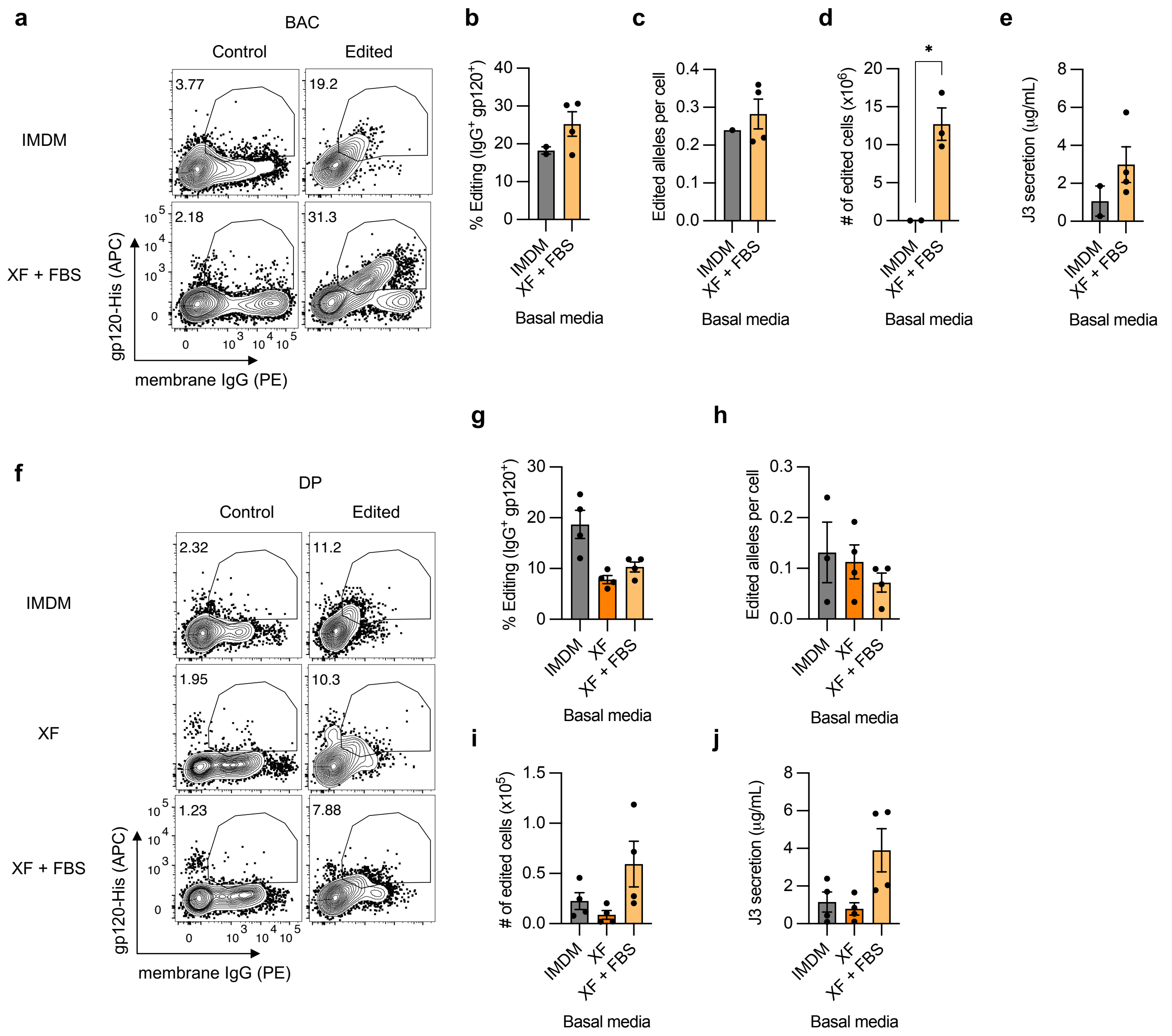


**Supplementary Figure 6. Editing in primary human B cells with different culture conditions.** Primary human B cells from $n$ = 3-4 independent experiments were activated with BAC (**a-e**) or DP (**f-j**) protocols in indicated basal media, starting at day -3, and edited at day 0 with sg05 Cas9 RNPs and an AAV6-J3 homology donor (MOI = 5 x $10^5$ vg/cell). (**a-b, f-g**) Editing rates were measured at day 8 by flow cytometry for surface J3-BCR. (**c,h**) Editing was quantified by in-out ddPCR at day 8 and normalized per cell against a control reaction. (**d,i)** The yield of edited cells at day 8 was calculated from 5 x $10^5$ starting B cells. (**e,j**) J3 HCAb secretion was measured by gp120-IgG ELISA at day 8. Some analyses for some replicates could not be performed due to low cell viability. Error bars show mean ± SEM. Statistics were calculated by 2-tailed t-test (b-e) or 1-way ANOVA (g-j). * $p < 0.05$.

**Supplementary Figure 7**

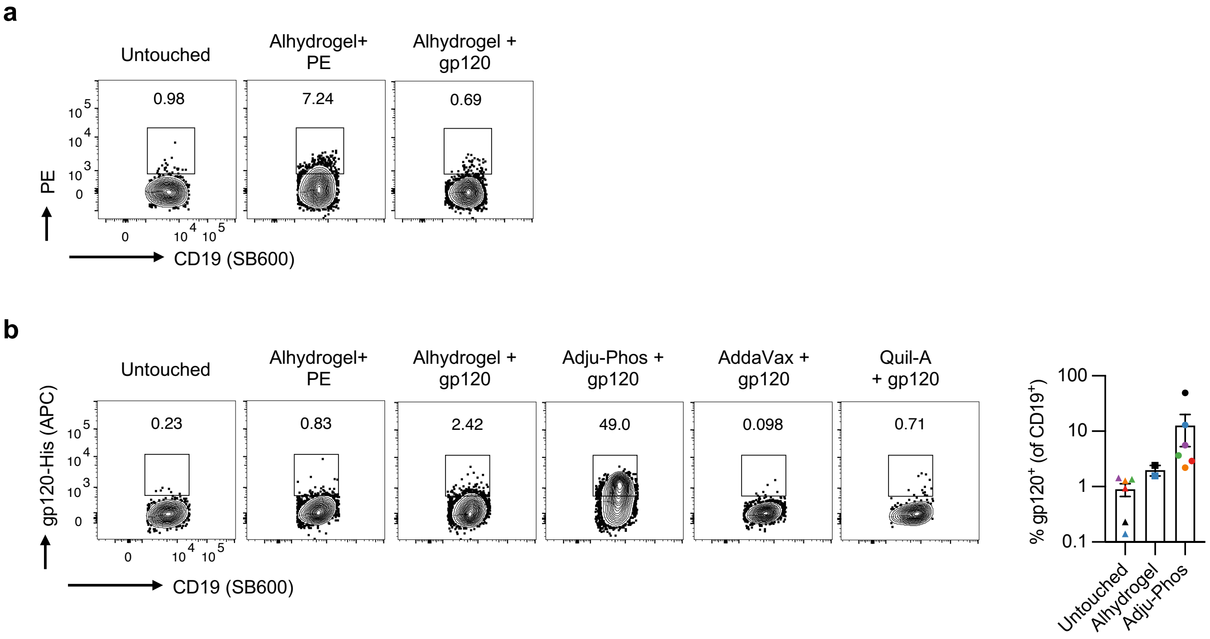


**Supplementary Figure 7. Tonsil organoid immunization.** (**a**) Antigen-specific response of tonsil organoids to immunization with Alhydrogel plus PE (as described in Wagar et al. 2021, *Nat Med* 27, 125-135). Antigen-specific responses were revealed by flow cytometry for PE-specific B cells 12 days after immunization, and gp120 immunization was included as a control. (**b**) Assessment of different adjuvants with gp120, by detection of gp120-specific B cells 12 days after immunization with gp120 or control PE antigens. Shown are representative plots from a single donor, and quantification of responses across donors (*n* = 2-6) comparing Alhydrogel and Adju-Phos adjuvants.

### Supplementary Figure 8

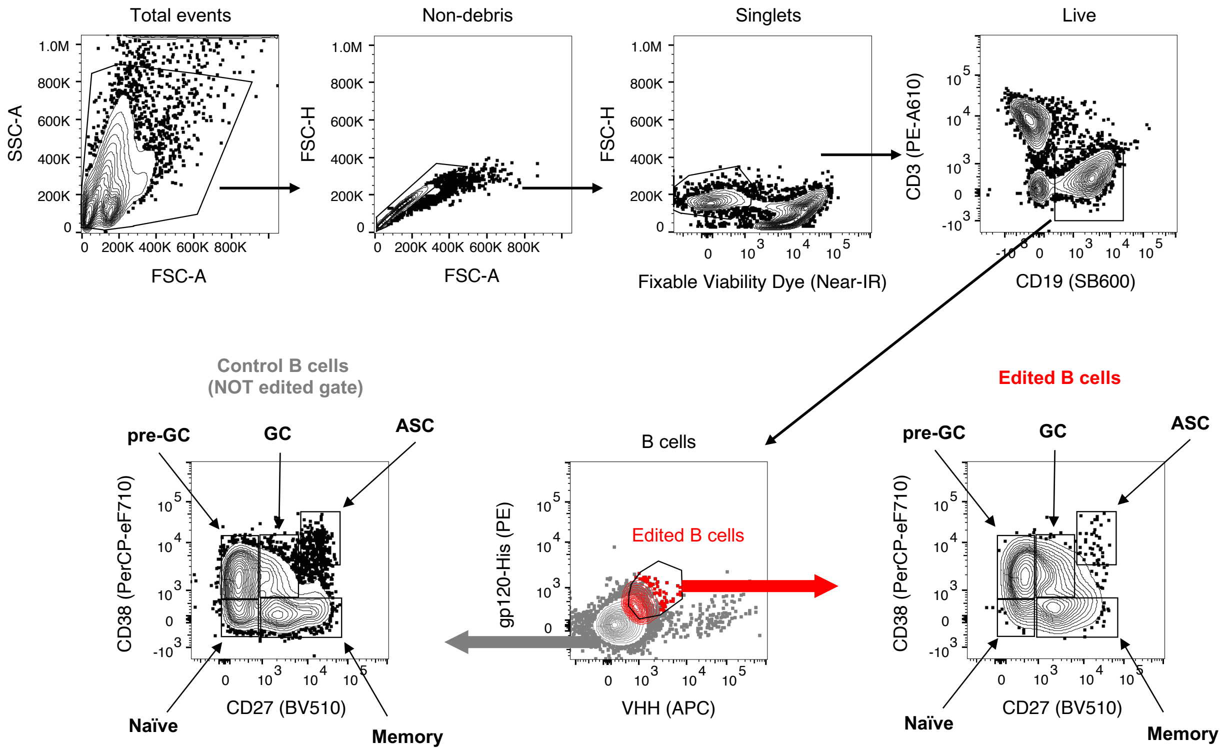


**Supplementary Figure 8. Representative gating for analysis of tonsillar B cells.** Tonsil organoids were generated by mixing total TMNCs and AAV6-J3 edited B cells in a 19:1 ratio, and optionally immunized with gp120 in Adju-Phos. Phenotypes of B cells were determined by flow cytometry after 12 days. Total lymphocytes were identified by excluding debris, non-singlets, dead cells, and on the basis of size, and B cells were identified as $CD19^+$ $CD3^-$. Edited B cells were gated as $V_HH^+$ $gp120^+$, and control B cells were the remainder of the B cells from the same sample not within the edited population gate. Naïve B cells were $CD27^-$ $CD38^-$, pre-germinal center (pre-GC) B cells were $CD27^-$ $CD38^+$, germinal center (GC) B cells were $CD27^+$ $CD38^+$, memory B cells were $CD27^+$ $CD38^-$, and antibody-secreting cells (ASCs) were $CD27^{hi}$ $CD38^{hi}$ (as described in Wagar et al. 2021, *Nat Med* 27, 125-135).
